# Dynamic Regulation of Innate Lymphoid Cell Development During Ontogeny

**DOI:** 10.1101/2023.11.01.565141

**Authors:** Tao Wu, Sijie Chen, Jie Ma, Xinyi Zhu, Maocai Luo, Yuanhao Wang, Yujie Tian, Qingqing Sun, Xiaohuan Guo, Jianhong Zhang, Xuegong Zhang, Yunping Zhu, Li Wu

## Abstract

Innate lymphoid cells (ILCs) are crucial for maintaining tissue homeostasis. The dynamic composition of ILC subsets during ontogeny has been observed for over a decade, yet the underlying mechanisms remain incompletely understood. Here, we combined differentiation assay and scRNA-seq analysis to compare the fetal and adult ILC development, and assessed their contribution to the dynamic change of ILC subsets. We revealed the preference for LTi cell differentiation in fetal hematopoietic progenitors, contrasted with a non-LTi differentiation bias in adult stages. Our analyses elucidated that the two stages shared a conserved differentiation trajectory and regulatory programs but used different controlling logic. Through computational prediction and dose-response experiments, we proposed a model where a transition from fetal RORγt-dominant states to adult Notch-GATA3-dominant states orchestrates these changes. This study suggests that ILC development can be modulated to accommodate the varying demands for specific ILC subsets within the body at different developmental stages.

**Highlight:**

- The ratio of LTi cells to non-LTi ILCs decreases from fetal to adult stage
- Modified fate determination of ILC progenitors contributes to this decrease of LTi cell ratio during ontogeny
- A conserved trajectory exists from early αLPs to Innate Lymphoid Cell Precursors (ILCPs) during both fetal and adult ILC development
- Enhanced activity of Notch-GATA3 signaling in ILC precursors accounts for the transition of their differentiation from LTi cells in the fetal to non-LTi ILCs in adults.

**Graphical abstract:** 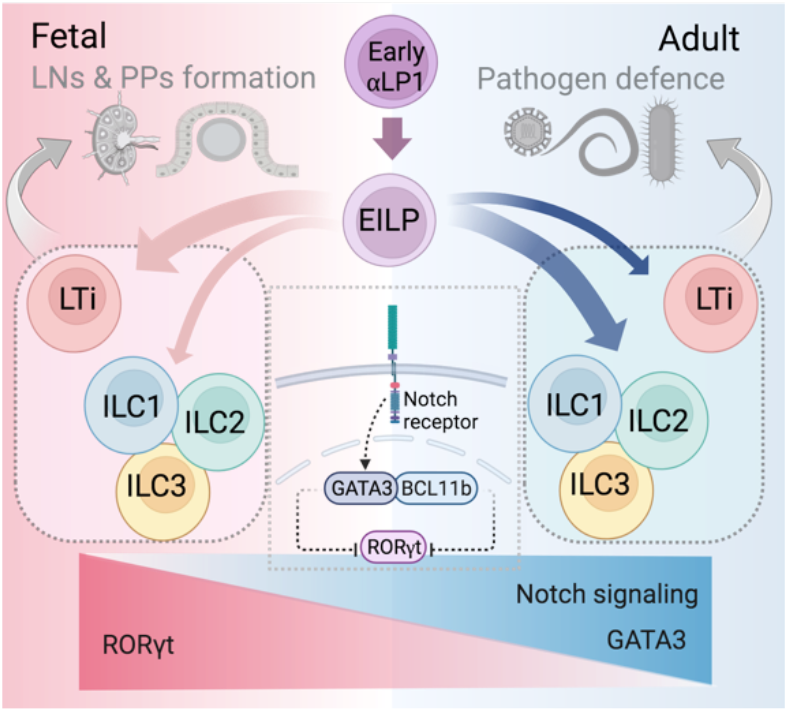

## INTRODUCTION

Innate lymphoid cells (ILCs), the innate counterparts of T cells (Vivier et al., 2018; Walker et al., 2013), encompass a diverse array of cell types including NK cells, ILC1s, ILC2s, ILC3s, and lymphoid tissue inducer (LTi) cells (Bal et al., 2020; Vivier et al., 2018). The ILC3s and LTi cells are also refered as RORγt^+^ ILCs. Originating predominantly from common lymphoid progenitors (CLPs), ILCs undergo differentiation through a series of progenitor stages, including early innate lymphoid progenitors (EILPs) (Yang et al., 2015), common helper innate lymphoid progenitors (CHILPs) (Klose et al., 2014), and innate lymphoid precursors (ILCPs) (Constantinides et al., 2014), all are included in α_4_β_7_^+^ lymphoid progenitors (αLPs). Different ILC subset pariticipate in different facets of immune responses: NK cells and ILC1s produce IFN-γ to defend against viruses and intracellular bacteria; ILC2s secrete IL-4/5/13, playing a role in defense against parasitic infections and in promoting allergies; ILC3s and LTi cells produce IL-17A/22 for defense against extracellular bacteria, crucial for maintaining local tissue homeostasis, particularly at mucosal sites (Bal et al., 2020; Vivier et al., 2018). LTi cells are also pivotal in forming secondary lymphoid tissues, such as lymph nodes and Peyer’s patches, primarily during the embryonic stage (Eberl et al., 2004; Onder and Ludewig, 2018).

Previous studies have demonstrated that the composition of ILC subsets exhibits dynamic shifts in various organs throughout murine development, reflecting adaptations to evolving physiological demands essential for maintaining homeostasis. LTi cells, predominant in the neonatal thymus, are scarcely present in adults, while ILC2s emerge as the dominant subset in the adult thymus (Jones et al., 2018). This trend of LTi cell proportions is also observed in the mouse small intestine (Sawa et al., 2010) and human tonsils and is associated with an increase in ILC3s (Hoorweg et al., 2012).

A consistent pattern in the shift of ILC subsets was suggested in the organs mentioned above. Nonetheless, the dynamic distribution of ILC subsets across various tissues remains to be systematically explored and summarized. Furthermore, while microenvironmental alterations are recognized as a primary external driver of ILC subset variations, the extent to which cell-intrinsic differentiation potential contributes to these variations remains ambiguous. Given the significant differences between hematopoietic progenitors at fetal and adult stages, these intrinsic cellular factors are likely to play a crucial role. Another significant challenge is the paucity of comprehensive profiling of ILC progenitors during fetal development, as current identification methods are primarily based on adult bone marrow studies.

In this study, we observed an elevated proportion of lymphoid tissue inducer (LTi) cells in various organs, including lymphoid (spleen) and non-lymphoid organs (small intestine, colon, lung, and liver) during fetal and newborn stages. Notably, the LTi cell ratio demonstrated a gradual decrease, concomitant with an increase in non-LTi innate lymphoid cells (ILCs) in adulthood. Concurrently, we found ILC progenitors from fetal liver exhibited a preferential differentiation towards LTi cells, whereas those derived from adult bone marrow showed an inclination to differentiate into non-LTi ILCs. This pronounced shift in differentiation preference was observed to occur within just one week post-birth. To elucidate the underlying mechanisms, we utilized single-cell RNA sequencing (scRNA-seq) to reconstruct the ILC developmental landscapes, identifying a conserved developmental trajectory from early αLPs to ILC precursors (ILCPs) in both fetal liver and adult bone marrow. Comparative analysis revealed enhanced Notch signaling and GATA3 expression, alongside reduced RORγt levels in adult progenitors compared to their fetal counterparts. Subsequent systematic dose-response experiments corroborated a “seesaw” model, articulating an interplay between Notch-GATA3 signaling and RORγt during murine ontogeny, which governs the dynamic shifts in ILC subsets. This model elucidates the predisposition of fetal ILC progenitors towards LTi cell differentiation, in contrast to the preference exhibited by adult progenitors.

## RESULTS

### Ratio of LTi cells to non-LTi ILCs decreases during ontogeny

To investigate the ontogenic distribution of ILCs, we systematically analyzed major ILC subsets in lymphoid (spleen) and non-lymphoid organs (small intestine, colon, lung, and liver) across various developmental stages in mice. The dynamic composition of ILC subsets in these organs reached a stable state within two weeks after birth (Figure 1 and Figure S1). Moreover, three categories of the ILC composition could be grouped based on their dominant subtype in adulthood (week 8): the ILC3-dominant category (small intestine in Figure 1A and 1B), the ILC2-dominant category(colon in Figure 1D and 1E, and lung in Figure S1A and S1B), and the ILC1-dominant category (spleen in Figure 1G and 1H, and liver in Figure S1D and S1E).

**Figure 1.**
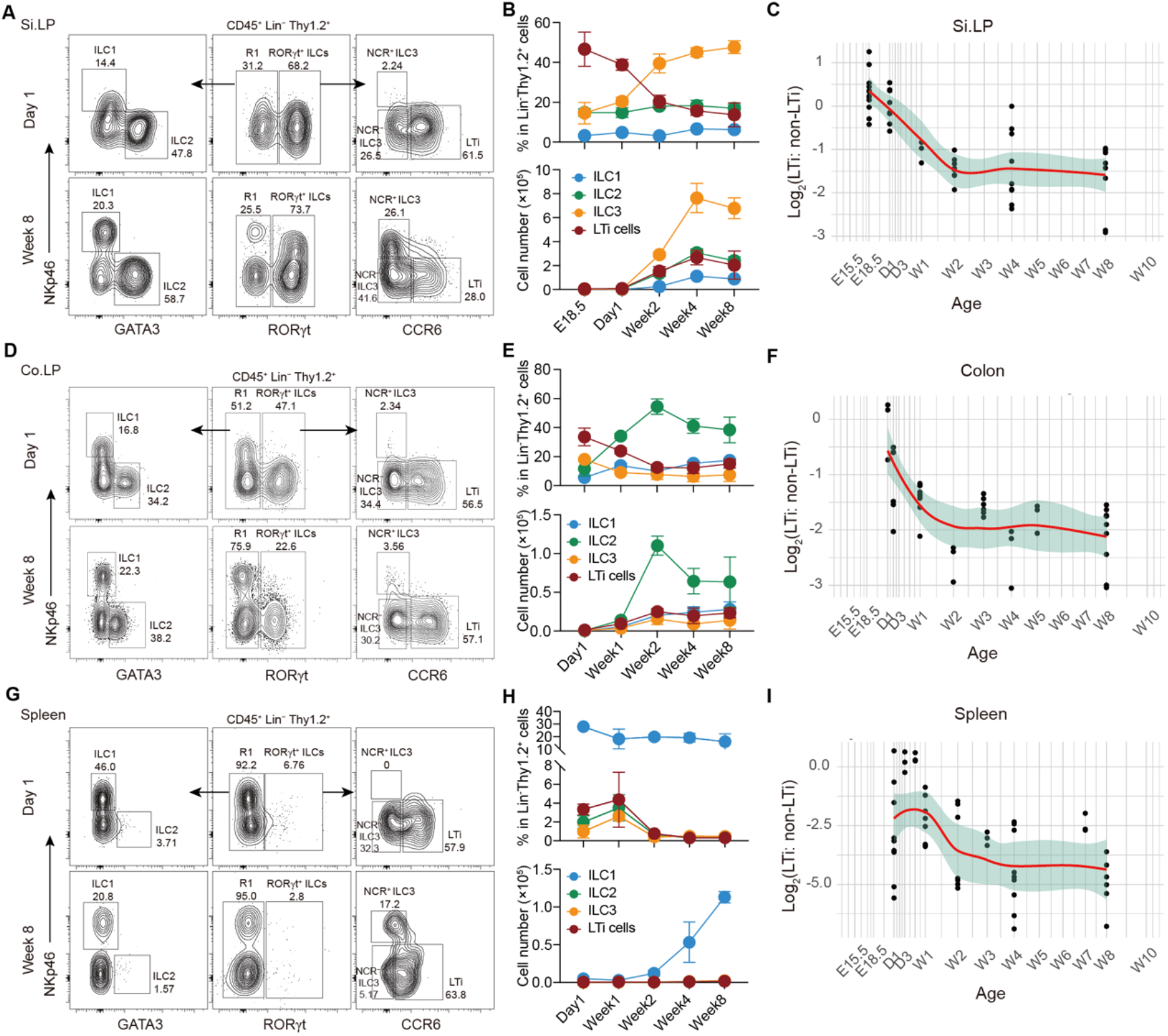
Temporal and tissue distribution of innate lymphoid cells during ontogeny. (A, D and G) Representative FACS plots show the frequencies of major ILC subsets in (A) lamina propria of small intestine (Si.LP), (D) lamina propria of colon (Co.LP) and (G) spleen, between newborn (Day 1) and adult (Week 8) mice. (B, E and H) The percentage and absolute number of ILC subsets in (B) Si.LP, (E) Co.LP and (H) spleen during ontogeny are shown. Data (n=3-6 mice per group) represent two independent experiments (mean ± SD). (C, F, I) The logarithm of the percentage ratio of LTi cells to non-LTi ILCs in (C) Si.LP, (F) Co.LP, and (I) spleen are plotted. Each dot represents the value for one individual mouse.

Despite the organ-specific variations in ILC subset distribution, a consistent pattern emerged, marked by a higher percentage of lymphoid tissue inducer (LTi) cells during the fetal and neonatal stages, transitioning to an elevated abundance of non-LTi ILCs, i.e., ILC1s, ILC2s, and ILC3s, in adulthood (Figure 1B, 1E, and 1H, Figure S1B and S1E). Concurrently, the ratio of LTi cells to non-LTi ILCs exhibited a gradual decline across all examined organs, reaching a stable state around two weeks postnatally (Figure 1C, 1F, and 1I, Figure S1C and S1F). This consistent shift suggested a potential shared mechanism governing the dynamic alterations in ILC subsets throughout ontogeny.

### Distinct progenitor differentiation capacity contributes to the composition shift of ILC subsets during ontogeny

While analysing ILCs, we also observed dynamic changes in αβT and γδT cells during ontogeny (Figure S2). Given the disparate capacities of hematopoietic progenitors to generate γδ T cells and αβ T cells during fetal and adult stages (Yuan et al., 2012), coupled with the close relationship between ILCs and T cells (De Obaldia and Bhandoola, 2015), we speculated that fetal and adult hematopoietic progenitors may exhibit differing abilities in generating LTi cells and non-LTi cells.

To investigate this hypothesis, we sorted hematopoietic progenitor cells (LSKs, CD45^+^Lin^−^c-Kit^+^Sca-1^+^) from fetal liver and adult bone marrow for an *in vivo* ILC differentiation assay. The resultant progenies in the small intestines of recipient mice were analyzed eight weeks later. Consistent with previous findings (Yuan et al., 2012), we observed a higher proportion of αβ T cells among the progenies derived from adult LSKs (Figure S3A). In contrast, the fetal LSK-derived cells exhibited higher proportions of ILCs and B cells (Figure S3A). Although both embryonic and adult LSKs exhibited proficiency in reconstituting all major ILC subsets (Figure S3B), a lower proportion of LTi cells and a higher abundance of ILC1s and ILC2s were identified among the ILCs derived from adult LSKs (Figure S3B and S3C), denoting an increased prevalence of non-LTi ILCs (Figure S3D). This observation aligned with the gradual reduction in the ratio of LTi cells to non-LTi ILCs throughout ontogeny.

To further compare the differentiation capacities of fetal and adult ILC progenitors, LSKs, CLPs, and αLPs were isolated from fetal liver and adult bone marrow for an *in vitro* differentiation assay. As expected, fetal ILC progenitors showed a predisposition for generating LTi cells (Figure S4A and S4C), whereas their adult counterparts leaned towards producing ILC1s and ILC2s, representing the non-LTi ILCs (Figure S4B and S4D). Furthermore, both fetal and adult αLPs exhibited enhanced efficiency in ILC differentiation compared to LSKs and CLPs (Figure S4C and S4D). Notably, a distinct population of NCR^+^ ILC3s was observed among the RORγt^+^ ILCs derived from adult αLP2s, whereas minimal to no NCR^+^ ILC3s was generated by fetal αLP2s, during both *in vitro* and *in vivo* differentiation assay (Figure S5). This divergence in ILC composition derived from fetal and adult αLP2s precisely mirrored the distinct distributions of ILC subsets between fetal and adult tissues.

Further analysis affirmed the consistent frequencies of CLPs, αLP1s, and αLP2s throughout different ontogenic stages (Figure 2A and 2B). To identify when the differentiation capacity begins to change, we sorted αLP2s from fetal liver and bone marrow at different ages for an *in vitro* differentiation assay. Remarkably, bone marrow αLP2s from mice within the first week post-birth exhibited significant changes in differentiation ability, characterized by a substantial decrease in the number of LTi cells and a simultaneous increase in non-LTi ILCs, akin to adult αLP2s (Figure 2C to 2F). This observation suggested that alterations in the differentiation capacity of ILC precursors occurred shortly after birth, approximately one week prior to the establishment of a steady-state in ILC subset composition across various tissues (Figure 1 and Figure S1). Consequently, the postnatal shift in the differentiation ability of ILC precursors could contribute to the dynamic changes of ILC subsets from the fetal to adult stages.

**Figure 2.**
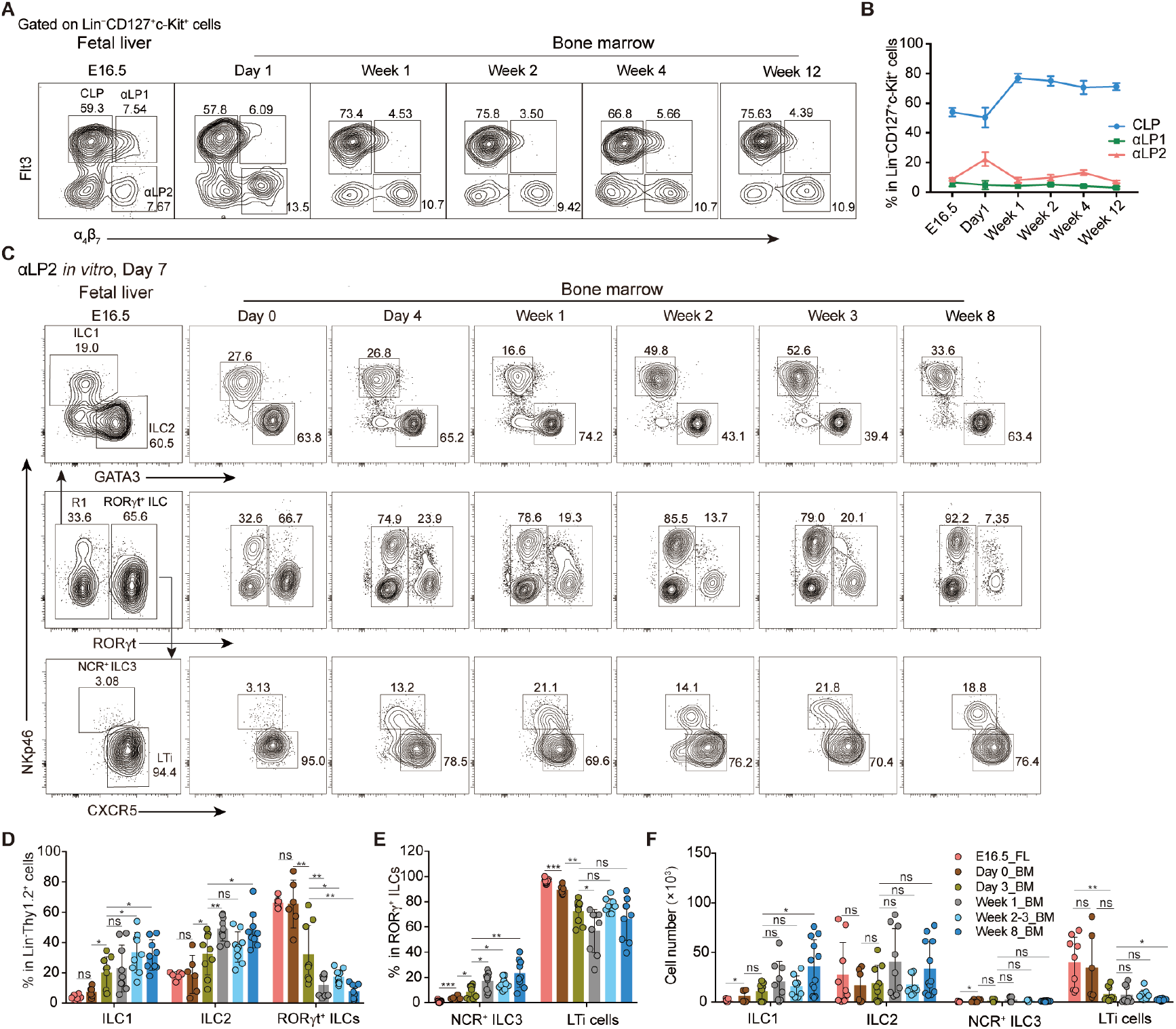
ILC subsets generated by progenitors from different stages during ontogeny. (A) CD45^+^Lin^−^CD127^+^c-Kit^int/+^ cells were gated to identify CLP, αLP1, and αLP2 in fetal liver (E16.5) and bone marrow at neonatal (Day 1), infant (Day 4, Week 1, and Week 2), weaning (Week 4), and adult stages (Week 12). (B) Dynamic percentages of CLPs, αLP1s, and αLP2s during ontogeny. Data (n=10 mice per group) are pooled from two independent experiments (mean ± SD). (C) Representative FACS plots show ILC subsets derived from αLP2s sorted from fetal liver (E16.5), or bone marrow at the newborn (Day 0), infant (Day 4 and Week 1), weaning (Week 2-3) and adult mice (Week 8), after 7 days of *in vitro* culture with IL-7 and SCF. (D-F) Percentage (D-E) and absolute cell number (F) of major ILC subsets in (C). Data (n=6-15 per group) are pooled from three independent experiments (mean ± SD).

### Single-cell transcriptomic landscape of fetal and adult ILC progenitors

ILC progenitors, rare populations in both fetal liver and adult bone marrow, have not been thoroughly elucidated, particularly in fetal tissues. To address this gap, we isolated αLPs, comprising both αLP1s and αLP2s, from fetal liver and adult bone marrow for single-cell RNA sequencing (scRNA-seq) analysis. Following stringent quality control procedures (see Methods), we selected 4284 fetal and 3442 adult αLPs for subsequent UMAP analysis. Ten distinct clusters were revealed (Figure S6A), all expressing of *Ptprc*, *Il7r*, *Itga4*, and *Itgb7* (Figure S6B). Clusters 0, 1, 2, and over half of cluster 6 exhibited high expression of *Flt3*, representing αLP1s. Conversely, clusters not expressing *Flt3* represented αLP2s (Figure S6B).

Based on the expression characteristics of known transcription factors, such as *Id2*, *Tcf7*, and *Zbtb16*, we defined clusters associated with the ILC lineage, specifically Cluster 6, 3, 7, 8, 9, and 4, primarily distributed within αLP2s (Figure S6B). Notably, Clusters 6, 3, and 9 represented early-stage progenitors in ILC development, evidenced by their high expression of *Tcf7*. A comparative mapping analysis of cell transcriptome (Methods) revealed that these early precursor cells shared some characteristics with early T cell progenitors (ETP) and double-ngative (DN) T cells (Figure S6C). Among them, Cluster 6 expressed high *Nfil3* and low *Id2*, representing early ILC progenitors (EILPs) (Harly et al., 2019). Cluster 3 showed high expression of *Zbtb16* and *Pdcd1*, serving as specific markers for ILC progenitors (ILCPs) (Constantinides et al., 2014; Yu et al., 2016). Additionally, Cluster 3 expressed signature genes of multiple ILC subsets, including *Gata3*, *Rora*, *Ccl5*, *Ltb*, etc., leading us to define it as ILCPs (Figure 3B). Cluster 7 also exhibited features of multiple ILC subsets, and high *Il18r1* expression, indicative of ILCPs seeded in the peripheral tissues via circulation system (Zeis et al., 2020), suggesting that Cluster 7 could represent a later stage of ILCPs. Accordingly, we labeled Clusters 3 and 7 as ILCP.1 and ILCP.2, respectively. Cluster 9, distinguished by high expression of *Rorc*, *Cxcr5*, *Ccr6*, and *Ltb*, represented LTi precursors. Additionally, Cluster 9 highly expressed *Cd74*, *H2-Aa*, and *H2-Ab1*, possibly associated with the newly annotated MHC-II^+^ LTi cells (Hepworth et al., 2013; Lyu et al., 2022). Meanwhile, Cluster 4 exhibited high features of both LTi cells (*Ltb*, *Rorc*, etc.) and ILC2s (*Gata3*, *Rora*, etc.) (Figure 3B), as confirmed by the cell-referring analysis (Figure S6C). Since Cluster 4 showed low expression of *Tcf7* and *Il18r1* (Figure 3B), it might be a mixed cluster containing LTiPs and ILC2Ps at a more terminal stage. Consequently, we defined Cluster 9 as an early stage LTiP.1 and Cluster 4 as LTiP.2/ILC2P. Finally, Cluster 8 was defined as NKPs based on their higher expression of *Eomes*, *Tbx21*, *Il2rb*, etc. (Figure 3B and S6C).

**Figure 3.**
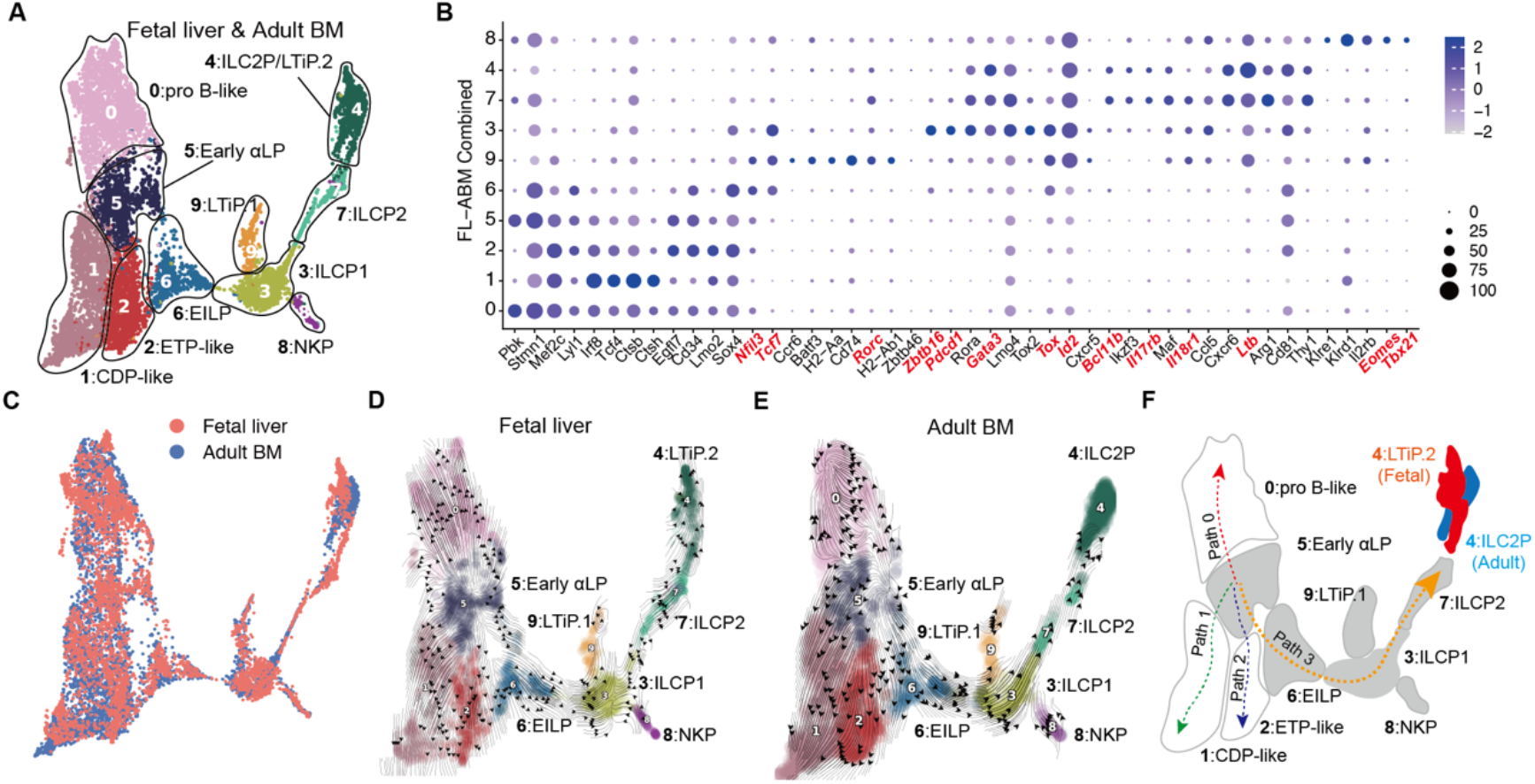
ILC differentiation trajectories in fetal liver and adult bone marrow revealed by single-cell RNA-sequencing analysis. (A) The cellular composition of αLPs from integrated fetal liver and adult bone marrow is visualized using Uniform Manifold Approximation and Projection (UMAP). Cell types are color-coded. (B) Feature gene expression is shown across the identified clusters among αLPs using dot-plot analyses. (C) The integrated UMAP shows αLPs from fetal liver (red) and adult bone marrow (blue). (D and E) RNA velocity algorithm-derived transition probabilities projected onto UMAP plots of (D) fetal liver and (E) adult bone marrow αLPs. Streamlines represent averaged and projected RNA velocities. (F) Schematic diagram showing the potential ILC differentiation trajectories in fetal liver and adult bone marrow.

Conversely, cells within Clusters 0, 1, 2, and 5, mainly comprising αLP1s, did not exhibit significant genetic features of the ILC lineage. Cluster 0 showed high expression of *Pbk* (Figure 3B), and a comparable fraction of Cluster 0 is transcriptionally similar to pro-B cells (Figure S6C). Therefore, we defined Cluster 0 as pro-B-like αLPs. Using the same strategy, we defined Cluster 1 as CDP-like αLPs due to their high expression of *Irf8*, *Tcf4*, *Ctsh*, *Ctsb*, *Siglech*, and Cluster 2 as ETP-like αLPs due to their high expression of *Cd34*, *Lmo2*, *Egfl7* (Figure 3B and S6C). Cluster 5 exhibited a high degree of heterogeneity, containing various cell types, and was labeled as early αLPs due to its early developmental characteristics (Figure 3B and S6C). Cluster 5 exhibited high expression of genes shared by Cluster 0, 1, 2, and 6, and contained multiple cell types, representing the highest heterogeneity among all these clusters according to cell referring analysis (Figure 3B and S6C). Therefore, we define Cluster 5 as the early αLPs.

Among the ten identified clusters, we observed an even distribution of cells from fetal liver and adult bone marrow, except for Cluster 4 (LTiP.2/ILC2P) (Figure 3C). Notably, fetal Cluster 4 was annotated as LTi cells, while adult Cluster 4 is annotated as ILC2, based on cell referring analysis (Figure S6D). Furthermore, fetal Cluster 4 showed higher expression of *Rorc*, *Cxcr5*, *Ccr6*, etc., indicating the LTiP moiety (Figure S6E), whereas adult Cluster 4 exhibited higher expression of *Rora*, *Gata3*, *Bcl11b*, *Il17rb*, etc., consistent with ILC2P characteristics (Figure S6G). Consequently, we refined the annotation, defining fetal Cluster 4 as LTiP.2 (Figure S6F) and adult Cluster 4 as ILC2P (Figure S6H).

### RNA velocity analysis predictes a conserved ILC differentiation trajectory in fetal liver and adult bone marrow

The markedly different capacity of fetal and adult ILC progenitors implicated potential distinct developmental pathways underlie these two stages. To further investigate this hypothesis, we employed RNA velocity analysis (Methods) to predict differentiation trajectories in both fetal and adult αLPs.

Four relatively conserved differentiation trajectories were predicted for both fetal and adult αLPs, all originating at early αLP (C5) and branching towards pro-B-like αLP (Cluster 0, Path 0), CDP-like αLP (Cluster 1, Path 1), ETP-like αLP (Cluster 2, Path 2), and ILC lineage (Cluster 3, Path 3), respectively (Figure 3D to 3F). Notably, a conserved ILC lineage (Path 3), initiated from early αLP (C5) and extending through EILP (C6) and ILCP (C3), was identified in both fetal liver and adult bone marrow. However, this ILC lineage (Path 3) terminated as LTiP in fetal liver, while transitioning to ILC2P in adult bone marrow (Figure 3D to 3F). This observation suggested that crucial regulatory factors may have undergone alterations from fetal to adult ILC development, resulting in distinct progenies of ILC progenitors despite the apparently conserved developmental pathway.

### RORγt- and GATA3-biased gene programs in fetal and adult ILC subset fates

To further explore the gene regulatory networks governing fetal and adult ILC development, we identified activated gene modules by applying the TSC (temporal spatial consensus) algorithm as described in the Method sections. We selected representative cells from the four trajectories (Figure 4A), analyzed the gene expression dynamics of these cells along Path 3, which represents the ILC commitment path from C5 to C3 (Figure 4B), and proposed a set of key gene regulatory modules along the path (Figure S7). Consistent with the conserved trajectories in both fetal and adult ILC development, we observed common patterns in gene expression dynamics. For example, we found that genes including *Flt3*, *Spi1*, *Irf8*, and *Lmo2* were notably enriched in the early stage of pseudotime, showing a significant decrease as differentiation progressed (Figure 4B). Following this, the induction of *Lck*, *Sox4*, *Tcf7*, and *Gfi1b* expressions marked the initiation of ILC commitment from early progenitor cells. This initiation was accompanied by the following expression of *Tox*, *Id2*, and *Zbtb16*, which further restricted the ILC commitment in these progenitors (Figure 4B).

**Figure 4.**
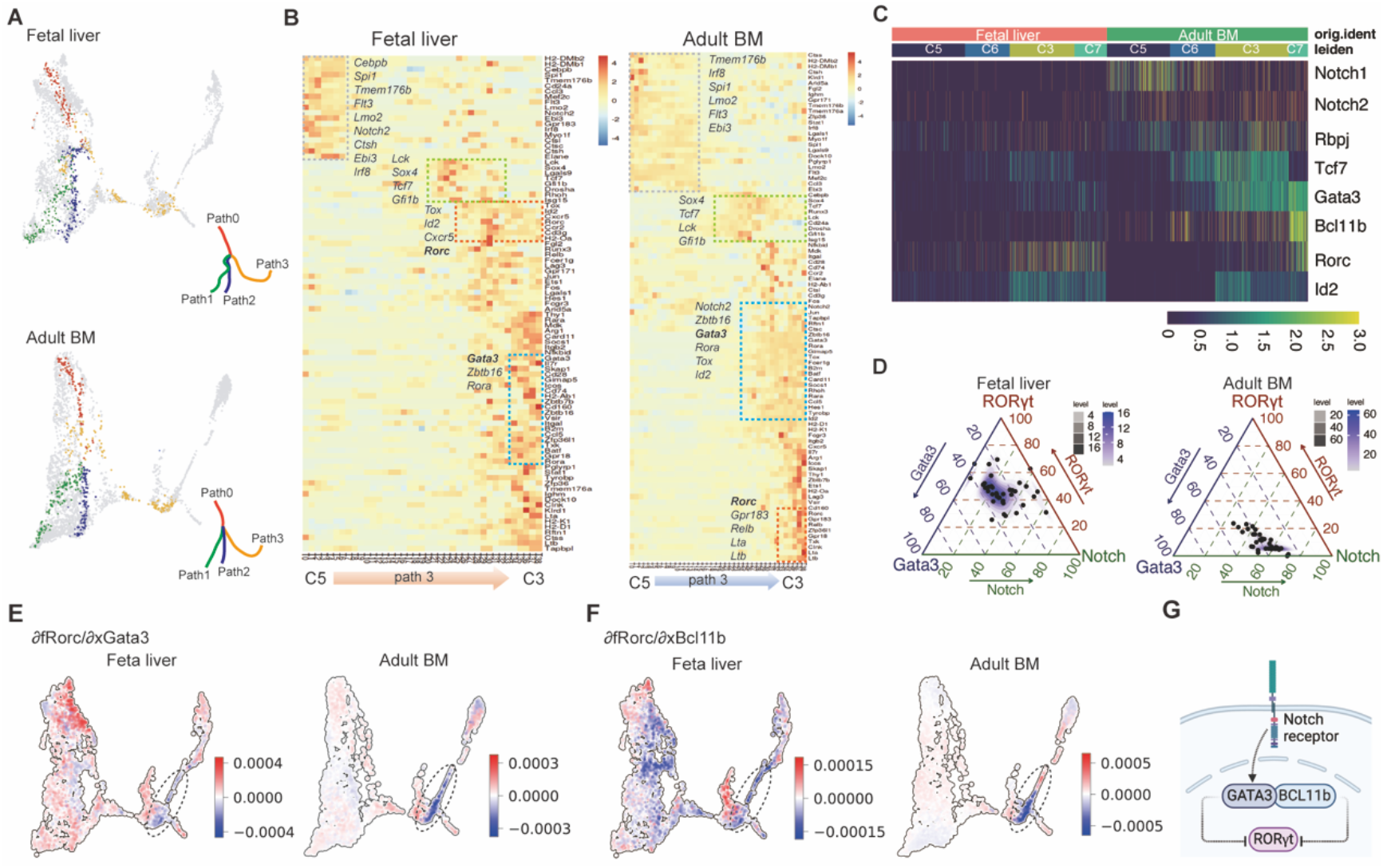
Comparation of gene programs during fetal and adult ILC development. (A) Visualization of the representative cells along the paths revealed in fetal liver and adult bone marrow αLPs. (B) Heatmaps displaying differentially expressed genes along the ILC differentiation pseudotime (path 3) from early αLP1s (C5) to ILCPs (C3) in fetal liver and adult bone marrow. (C) Heatmap of gene expressions associated with Notch signaling, *Gata3*, *Bcl11b*, and *Rorc* during both fetal and adult ILC differentiation. (D) Ternay plots showing mutual relationships among activation scores of Notch signaling pathway, GATA3 regulon, RORγt regulon. The activation scores were evaluated along the ILC differentiation trajectories (Path 3) in fetal and adult stages. Colors of the dashed lines indicate the projection direction of the coordinates. (E and F) *Dynamo* analysis predicting the potential relationship between (E) *Rorc* and *Gata3*, as well as between (F) *Rorc* and *Bcl11b* during fetal and adult ILC development trajectories. The dashed circles highlighted the negative regulatory relationships revealed by negative Jacobian values. (G) Schematic diagram showing the potential relationship among Notch signaling, GATA3/BCL11b and RORγt.

Nevertheless, the gene expression programs had exhibited major different logic between the fetal and adult stages. Specifically, in the fetal stage, we observed that the expression of *Rorc* was nearly synchronous with *Tox* and *Id2*, whereas the *Gata3* expression was relatively delayed (Figure 4B and S7A). Conversely, in the adult stage, the *Gata3* expression closely paralleled that of *Tox*, *Id2*, and *Zbtb16*, whereas the peak expression of *Rorc* lagged behind (Figure 4B and S7B). Moreover, our analysis of differentially expressed genes revealed elevated expression levels of *Gata3* and *Bcl11b* in the adult clusters, particularly in C6 (EILP) and C3 (ILCP), but higher *Id2* and *Rorc* in the fetal clusters (Figure S8A). Additionally, higher GATA3 expression in adult ILCPs (C3: GATA3^hi^TCF-1^+^ αLP2) and lower RORγt in the adult clusters (C9: RORγt^hi^ TCF-1^+^ αLP2) were confirmed by the flow cytometry analyses (Figure S8D to S8F). Consistent with these findings, the expression level of RORγt in intestinal RORγt^+^ ILCs gradually decreased during the ontogeny (Figure S8G and S8H).

Notch signaling plays a crucial role in ILC differentiation, especially the ILC2s and ILC3s (Possot et al., 2011; Wong et al., 2012). Notably, we observed that *Notch2* was transiently expressed in the early stage of fetal ILC lineage (Path 3), followed with a significant downregulation. Conversely, in the adult stage, *Notch2* expression nearly synchronized with *Zbtb16* and *Gata3* (Figure 4B). Moreover, *Rbpj*, a pivotal downstream component of the Notch receptor, was also synchronously expressed with *Tox* and *Id2* during adult ILC commitment (Figure S7B), in contrast to its limited temporal expression during fetal ILC commitment (Figure S7A). Our analyses of differentially expressed genes uncovered elevated expression levels of *Notch1* and *Rbpj* in adult clusters (Figure S8A). Additionally, higher Notch2 expression in adult αLPs was confirmed by flow cytometry analysis (Figure S8B and S8C). Gene Set Variation Analysis (GSVA) further indicated a notably enriched Notch signaling pathway in adult αLPs (Figure S9). Consequently, a heightened level of Notch signaling activity was discerned during ILC development in the adult stage than in the fetal stage.

The induction of GATA3 and Bcl11b as a consequence of Notch signaling activation has been documented in previous studies (Ho et al., 2009; Hosokawa and Rothenberg, 2021; Rothenberg, 2019). These factors are known to exert repression on *Rorc* expression (Califano et al., 2015; Zhong et al., 2015). During adult ILC commitment, we observed an elevation in the expression of *Gata3* and *Bcl11b* (C6 and C3), closely following the heightened expression of *Notch1*, *Notch2*, and *Rbpj* (C5), key components of the Notch signaling pathway. This trend was less apparent during fetal ILC development (Figure 4C). Additionally, an evident opposing expression pattern between *Gata3*/*Bcl11b* and *Rorc* became apparent. Elevated *Gata3* and *Bcl11b* expression aligned with decreased *Rorc* expression during adult ILC commitment. Conversely, elevated *Rorc* expression was associated with decreased *Gata3* and *Bcl11b* during fetal ILC commitment (Figure 4C).

Furthermore, we quantified the activation levels of the Notch signaling pathway, the GATA3 regulon, and the RORγt regulon during ILC differentiation as well as their mutual regulations (Method). We employed ternary plots to analyze the interplay between them, as depicted in Figure 4D. The cell states (points on the ternary plots) shifted from a RORγt-dominant distribution (fetal) to a Gata3-Notch dominant distribution (adult). These data suggested that RORγt may act as a key transcription factor during fetal ILC development, whereas Notch signaling and GATA3 appear to be more influential in modulating adult ILC development. With the help of *dynamo* analysis (Qiu et al., 2022), we further predicted a suppressive effect of GATA3 and Bcl11b on *Rorc* expression, particularly in the adult ILCP (C3) (Figure 4E and 4F). The analysis calculated Jacobian values for each cell and reported negative partial derivatives in the ILCP zones on the UMAP.

Based on the above observations and previously reported discoveries, we inferred a regulatory model that explains the ILC subset preferences during fetal and adult stages. In this model, we suggested that the amplified Notch signaling observed in adulthood likely leads to an upregulation of GATA3 and Bcl11b. This in turn results in the downregulation of Rorc expression, as detailed in Figure 4G.

### Enhanced activity of Notch-GATA3-signaling suppresses the generation of LTi cells but promotes non-LTi ILCs

Given the pivotal roles of Notch signaling, GATA3, and RORγt in the developmental process of ILCs (Chea et al., 2016b; Cherrier et al., 2012; Eberl et al., 2004; Possot et al., 2011; Sanos et al., 2009; Vonarbourg et al., 2010; Yagi et al., 2014; Zhong et al., 2015; Zhong et al., 2020), we speculated that changed Notch signaling, GATA3 and RORγt expression may result in the modified fate determination of ILC subsets from the fetal to the adult stage.

To address this question, we performed an *in vitro* differentiation assay of ILCs using the OP9-DL1 cell line as feeder cells, which provides continuous ligands for activation of Notch signaling. Our experiments found that sustained activation of Notch signaling led to a substantial increase in the proportion of ILC2s (Figure 5A), which was consistent with prior studies (Wong et al., 2012). The ILC2s increase was acompanied by a significant reduction in the proportion of LTi cells among the progenies of fetal ILC progenitors (Figure 5B) and a notable decrease in the proportion of ILC1s among the progenies of adult ILC progenitors (Figure 5C). The proportion of ILC3s exhibited no significant change during this process (Figure 5D). Thus, enhanced Notch signaling promoted the generation of ILC2 while suppressing LTi cell development. This phenomenon could explain why the ILC lineage in the fetal liver ended with LTiPs, while in adult bone marrow with ILC2Ps (Figure 3D to 3F). Moreover, intervening in Notch signaling by DAPT at early stages, such as CLP and αLP1, resulted in a more substantial alteration of ILC subsets (Figure 5A to 5D). Additionally, blocking Notch signaling in αLP2s could significantly reduced their ability to differentiate into ILC3s, leading to the decreased proportion of ILC3s and increased proportion of LTi cells (Figure S10). However, this intervention had limited effects on the generation of ILC1s and ILC2s (Figure S10).

**Figure 5.**
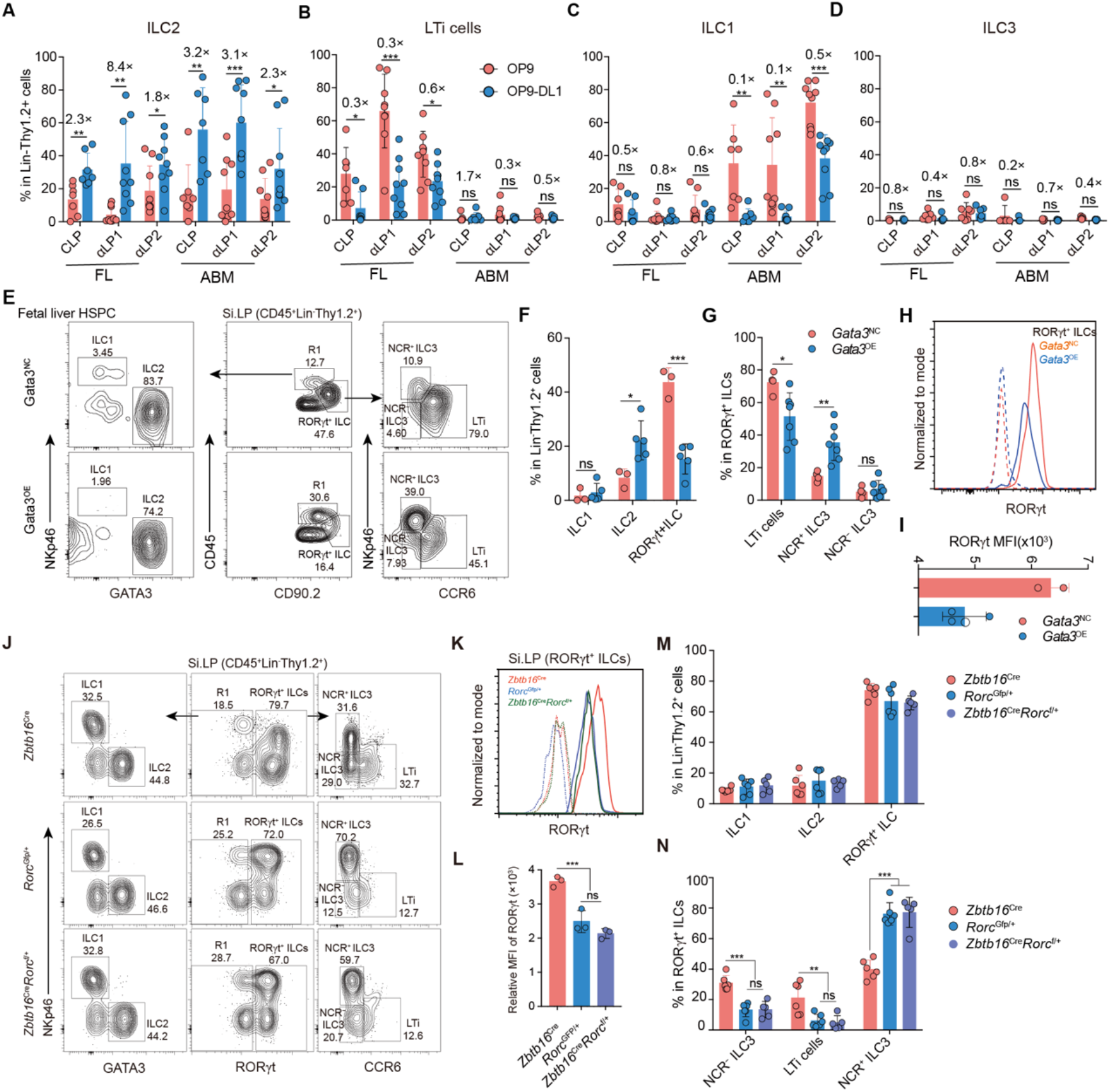
Enhanced Notch signaling, increased GATA3 and decreased RORγt suppresses LTi cell differentiation. (A-D) Percentage of ILC1s, ILC2s, ILC3 and LTi cells derived from fetal and adult CLPs, αLP1s and αLP2s after co-culture with either OP9 or OP9-DL1 stromal cells *in vitro*. Data (n=7-8 mice per group) are pooled from three to four independent experiments (mean ± SD). (E) Representative flow cytometry plots showing the ILC subsets derived *in vivo* from the fetal liver progenitors transduced with either empty plasmid (*Gata3*^NC^) or *Gata3* overexpression (*Gata3*^OE^) in the recipient intestine. (F-G) The percentage of ILC subsets in (E). Data (n=3-7 mice per group) are pooled from two independent experiments (mean ± SD). (H and I) Intracellular staining of RORγt in RORγt^+^ ILCs in (G). Data (n= 2-3 mice per group) represent two independent experiments (mean ± SD). (J) Flow cytometry analysis showing the major ILC subsets in the small intestine of *Rorc* heterozygous mice, including both *Rorc*^Gfp/+^ and *Zbtb16*^Cre^*Rorc*^f/+^ mice. (K and L) Mean fluorescence intensity (MFI) of RORγt in small intestinal RORγt^+^ ILCs from *Rorc* heterozygous mice. Data (n=3 per group) represent two independent experiments (mean ± SD). (M and N) Percentage of major ILC subsets in the small intestine from *Rorc* heterozygous mice. Data (n=3 per group) represent two independent experiments (mean ± SD).

To assess the impact of elevated GATA3 expression on ILC differentiation from fetal to adult stage, we conducted *Gata3* overexpression experiments in fetal hematopoietic stem/progenitor cells (HSPCs). Subsequently, these modified HSPCs were then transplanted into recipient mice to observe *in vivo* ILC differentiation. Analysis of the progenies derived from GATA3-overexpressing HSPCs revealed a significant increase in the porpportion of ILC2s, accompanied by a notable decrease in the proportion of total RORγt^+^ ILCs. Within the RORγt^+^ ILCs, Gata3 overexpression led to an increased proportion of NCR^+^ ILC3s and a decreased proportion of LTi cells (Figure 5E to 5G). Furthermore, *Gata3* overexpression resulted in a reduction in the expression level of RORγt in RORγt^+^ ILCs isolated from the recipient small intestine (Figure 5H and 5I). This observation aligned with the previously established negative regulatory effect of GATA3 on *Rorc* expression (Figure 4E to 4G).

The reduced RORγt expression, as shown in adult *Rorc* heterozygous mice with both *Rorc*^Gfp/+^ and *Zbtb16*^Cre^*Rorc*^f/+^ genotypes (Figure 5K and 5L), led to decreased proportion of LTi cells among RORγt^+^ILCs and substantially increased proportion of NCR^+^ILC3s (Figure 5J and 5N). However, the proportions of ILC1s and ILC2s showed no significant change in *Rorc* heterozygous mice (Figure 5J and 5M). Additionally, upon transplanting the LSKs from the adult *Rorc* heterozygous mice for *in vivo* ILC differentiation analysis, we found a dramatically decreased proportion of total RORγt^+^ILCs. Within this population, the proportion of LTi cells was significantly reduced, while the proportion of NCR^+^ILC3s showed a remarkable increase (Figure S11A and S11B).

Furthermore, we observed that NCR^+^ILC3s emerged three days after birth in the small intestine of *Rorc*^Gfp/+^ mice, a significantly earlier occurrence compared to wild-type mice (Figure S11C). This observation indicated an accelerated induction of NCR^+^ILC3s in mice when the RORγt expression level was halved. Notably, an increased proportion of the NCR^+^ILC3 population was also generated by fetal liver αLPs from *Rorc*^Gfp/+^ mice, compared to that derived from wild type fetal αLPs (Figure S11D and S11E). Additionally, an elevated proportion of NCR^+^ILC3s and a reduction in LTi cells were detected among the progenies of adult *Rorc* heterozygous αLP2s (Figure S11F and S11G). In contrast, upon further overexpressing RORγt in adult hematopoitic stem/progenitor cells (HSPCs), a decreased proportion of NCR^+^ILC3s and an increased proportion of LTi cells were observed among the progenies in the recipient small intestine, while the development of ILC1s and ILC2s remained largely unaffected (Figure S11H to S11L).

Taken together, these findings demonstrated a hierarchical molecular regulation of ILC differentiation throughout mouse ontogeny. Specifically, a higher level of RORγt expression by fetal ILC progenitors correlated with an increased potential to preferentially generate LTi cells, particularly during the fetal and neonatal stages. Conversely, enhanced Notch-GATA3 signaling in the ILC precursors during later developmental stages was associated with decreased RORγt expression, fostering the generation of non-LTi ILCs while suppressing LTi cell production. Consequently, the dynamic modulation in the fate of different ILC subsets during ontogeny appears to be driven by a seesaw-like transition: moving from a RORγt-dominant phase in fetal stages to a stage characterized by enhanced Notch signaling and GATA3 in adulthood.

## DISCUSSION

ILCs play a crucial role in maintaining tissue homeostasis within the immune system, with different subsets mediating distinct immune response (Bal et al., 2020; Vivier et al., 2018). Dynamic alterations in ILC subsets have been documented in the murine small intestine (Sawa et al., 2010) and thymus (Jones et al., 2018) in recent years. However, a comprehensive elucidation of this phenomenon across various tissues during ontogeny is yet to be achieved. In this study, we proposed a “seesaw” model of the interplay between Notch-GATA3 signaling and RORγt during murine ontogeny. Specifically, RORγt demonstrated enhanced regulatory influence in fetal ILC development, whereas the Notch signaling pathway and GATA3 assumed augmented regulatory functions in adulthood. This model elucidated the predisposition of fetal ILC progenitors towards LTi cell differentiation, contrasting with the preference of non-LTi ILCs exhibited by adult progenitors. This dynamic regulation of ILC progenitors may account for the evolving composition of ILCs during ontogeny.

It is reasonable that different ILC subsets are required during various developmental stages. Remarkably, our findings, along with those of others, consistently demonstrate a prevalence of LTi cells throughout fetal and newborn tissues, followed by an increased proportion of non-LTi ILCs, including ILC1s, ILC2s, and ILC3s. The significant role of LTi cells in the formation of secondary lymphoid organs (Eberl et al., 2004; Krishnamurty and Turley, 2020), particularly during the embryonic and newborn stages (Onder and Ludewig, 2018), suggests a prioritization in the body to establish key secondary lymphoid tissues in anticipation of the impending immune defense post-birth. However, with the direct exposure to various pathogens after birth, non-LTi ILCs become critical to defense against various pathogens (Eberl et al., 2015; Lim and Di Santo, 2019; Oherle et al., 2020; Vivier et al., 2018), particularly during the early stages after birth, when the T cell-mediated adaptive immune response is incompletely formed (Mao et al., 2018).

Recent studies have emphasized the impact of local niches on this process (Lim et al., 2017; Oherle et al., 2020; Zeis et al., 2020). For instance, a high concentration of AhR ligands in the small intestine promotes the differentiation and proliferation of ILC3s while restricting the expansion of ILC2s (Kiss et al., 2011; Lee et al., 2012; Li et al., 2018). In the lung environment, regardless of whether these precursor cells originate from neonatal or adult sources, they are driven to differentiate into ILC2s (Ghaedi et al., 2020; Oherle et al., 2020; Zeis et al., 2020). However, these findings could not fully explain the phenomenon observed in this study, where the ratio of LTi cells to non-LTi ILCs systematically decreased across tissues during ontogeny. Therefore, additional mechanisms regulating the dynamic changes of ILC subset composition require further clarification.

It is well-established that the characteristics of hematopoietic stem/progenitor cells (HSPCs) change during ontogeny (Jassinskaja et al., 2017; Kim et al., 2007; Medvinsky et al., 2011; Yuan et al., 2012). Fetal HSPCs exhibit a higher capacity of γδ T cells generation, while adult HSPCs show an enhanced capacity of αβ T cells generation (Yuan et al., 2012). Correspondingly, fetal tissues manifest a higher composition of γδ T cells, whereas adult tissues exhibit a higher proportion of αβ T cells (Dunon et al., 1997). In line with the widely shared regulatory mechanisms governing T cell and ILC development, our findings demonstrate distinct capacities for generating ILC subsets from fetal and adult hematopoietic progenitors. Particularly noteworthy is the higher capacity for LTi cell generation by fetal progenitors, contrasting with the greater capacity for non-LTi ILC generation by adult progenitors. This pattern aligns with the systematic reduction in the ratio of LTi cell to non-LTi ILC from the fetal to adult development. Therefore, we propose that hematopoietic progenitors, exhibiting distinct differentiation capacities for ILC subsets at different developmental stages, play a significant role in driving the dynamic changes in ILC subset composition during ontogeny.

A series of ILC progenitors has been identified in adult bone marrow (De Obaldia and Bhandoola, 2015; Harly et al., 2019; Ren et al., 2022; Vivier et al., 2018), however, there is still a lack of comprehensive profiling of ILC progenitors during fetal development (Ishizuka et al., 2016). Notably, the development of fetal and adult ILCs exhibits distinct sensitivities to the deficiency of Notch signaling (Chea et al., 2016a; Possot et al., 2011) and Ikaros (Li et al., 2016; Schjerven, 2013), implying potential different ILC developmental pathways between these two stages. Contrary to expectations, our scRNA-seq analyses revealed similar types of ILC progenitors and a conserved differentiation trajectory during fetal and adult ILC development. However, fetal and adult progenitors exhibited different transcriptomes and gene regulatory networks. Remarkably, Notch signaling was enriched during adult ILC development, as evidenced by higher expression of *Gata3*, *Tcf7*, and *Bcl11b*. These transcription factors, involved in ILC subset fate determination, are all known to be promoted by Notch signaling (Cherrier et al., 2018; Fang and Zhu, 2017). Moreover, TCF-1 (encoded by *Tcf7*) (Mielke et al., 2013; Yang et al., 2015), GATA3 (Zhong et al., 2016; Zhong et al., 2020) and Bcl11b (Califano et al., 2015) all show a higher requirement for non-LTi ILC generation, concurrently suppressing LTi cell differentiation. This aligns with the observation that adult ILC3s, not fetal LTi cells, depend on Notch signaling (Chea et al., 2016a; Possot et al., 2011). Consistent with prior observations that GATA3 and Bcl11b can suppress RORγt expression (Califano et al., 2015; Zhong et al., 2020), we aso confirmed reduced RORγt expression in the adult. Moreover, reduced RORγt expression accelerated the emergence of ILC3 and suppressed LTi cell generation. Therefore, we propose the modified activity of Notch signaling, leading to changed expression of Gata3 and Rorc, will altered the fate determination of ILC subsets and ultimately contribute to the dynamic change of ILC subsets during ontogeny.

Furthermore, Notch signaling also plays a role in determining the fate of αβ T cells and γδ T cells in a dose-dependent manner. Increased Notch signaling activity is necessary for the generation of αβ T cells, while it is not essential for γδ T cells (Ciofani et al., 2006; Garbe et al., 2006; Tanigaki et al., 2004). Taking into account the enhanced Notch signaling observed in adult ILC progenitors, which still maintain T cell differentiation capacity, exemplified by the early αLP1 cluster (C5) we identified, it is plausible that enhanced Notch signaling could serve as a common trigger for shifting differentiation capacity from LTi cells to non-LTi ILCs, and probably from γδ T cells to αβ T cells as well, during the transition from fetal to adult stages. However, the mechanisms underlying the modulation of Notch signaling strength during ontogeny remain unclear and warrant further investigation.

Collectively, our findinngs highlight the dynamic regulation of ILC subset fate determination by adjusting the activity of Notch-GATA3 signaling and RORγt expression, leading to the gradually decreased ratio of LTi cells to non-LTi ILCs from fetal to adult. This study propose an additional mechanism that altered differentiation capacity of ILC progenitors may coordinate with the environment cues to meet the demands of distinct ILC subsets in the body during mouse ontogeny.

## Limitations of Study

Our study revealed a correlation between changes in the differentiation ability of ILC progenitor cells and the dynamic composition of ILC subsets in tissues during ontogeny. However, direct evidence for the extent to which changes in differentiation ability contribute to the distribution of ILC subsets in peripheral tissues is lacking. While we demonstrated that the differentiation capacity of ILC progenitors changed as early as one week after birth, we were unable to determine definitively whether ILC progenitors from the bone marrow are still required for generating ILC subsets in peripheral tissues after birth. This consideration is particularly pertinent given the tissue-residence feature of ILCs in adults.

## AUTHOR CONTIBUTIONS

T.W., S.C., and L.W. designed the study. T.W. performed the experiments and acquired data with assistance of X.Zhu. S.C. analyzed the single-cell RNA-seq data. Y.Z., J.M., and J.Z. provide the single-cell RNA-seq data. M.L., Y.T., Q.S., X.G. and Y.W. provided technical and experimental assistance. T.W., S.C., L.W. analyzed and interpreted the data. Manuscript was written by T.W., S.C. and L.W., S.C. and J.M. are responsible for the data curation, and this study was supervised by L.W., Y.Z. and X.Zhang.

## ACKNOWLEDGEMENTS

We thank Xiaoguang Li, Wenlong Lai, Yujie Tian, Yijun Zhang, Jiaoyan Lv, Xin Liu, Chao Wang, Zen Shi, Site Feng, Haoxiang Gao, Ning Wu and Jingjiao Wu for for administrative/support. We thank Qinli Sun from Dr. Chen Dong lab in this institute for providing the *Rorc* over-expression plasmid. We thank Dr. Yan Shi, Dr. Xiaoyu Hu and Dr. Wenwen Zeng for providing advice. Flow cytometry and cell sorting were carried out on IITU platforms. We also thank the assistance of Laboratory Animal Research Center of Tsinghua University. This study was supported by grants from the Ministry of Science and Technology of China (National Key Research Project: 2019YFA0508502 and 2022YFC2505001 to L.W.; 2020YFE0202200 to Y.Z. and J.M.; 2021YFF1200900 to X.Z.), the National Natural Science Foundation of China (Grants 31991174 to L.W.; Grant 62250005 to X.Z.), the National Basic Research Center of China (82388101 to L.W.), China Postdoctoral Science Foundation (2022M721818 to T.W.) and Tsinghua-Peking Center for Life Sciences (to L.W. and T.W.).

## METHODS

### KEY RESOURCE TABLE

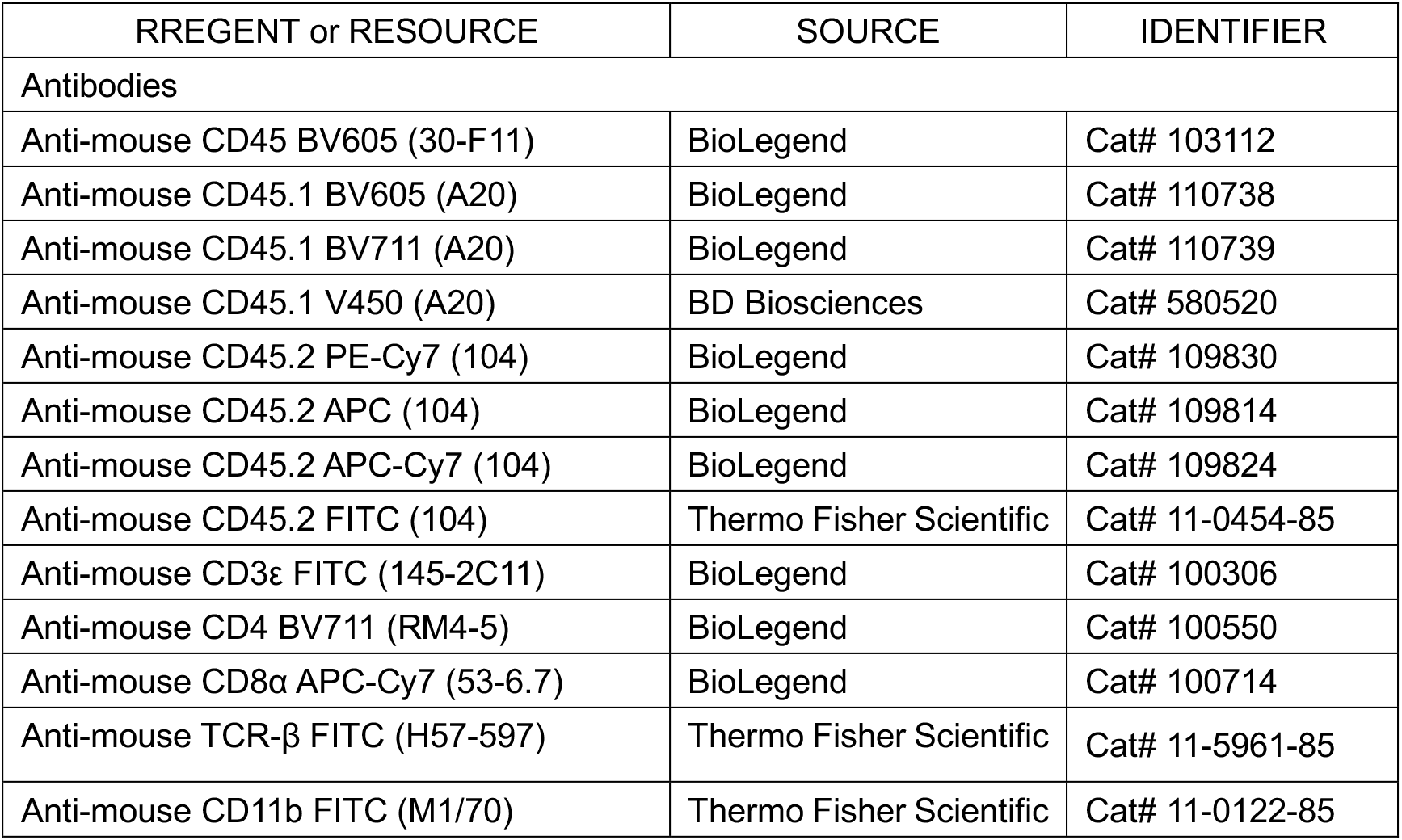
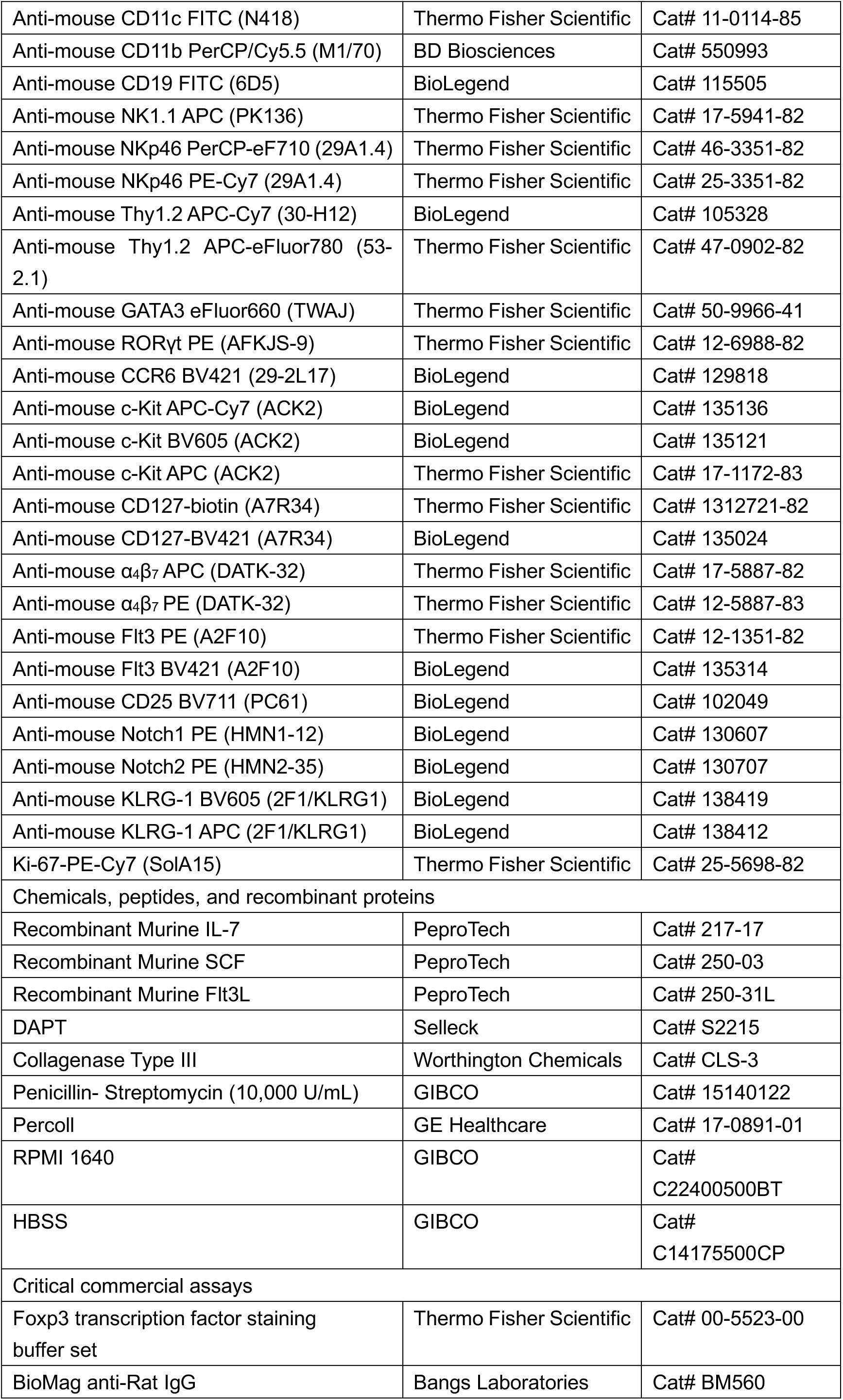
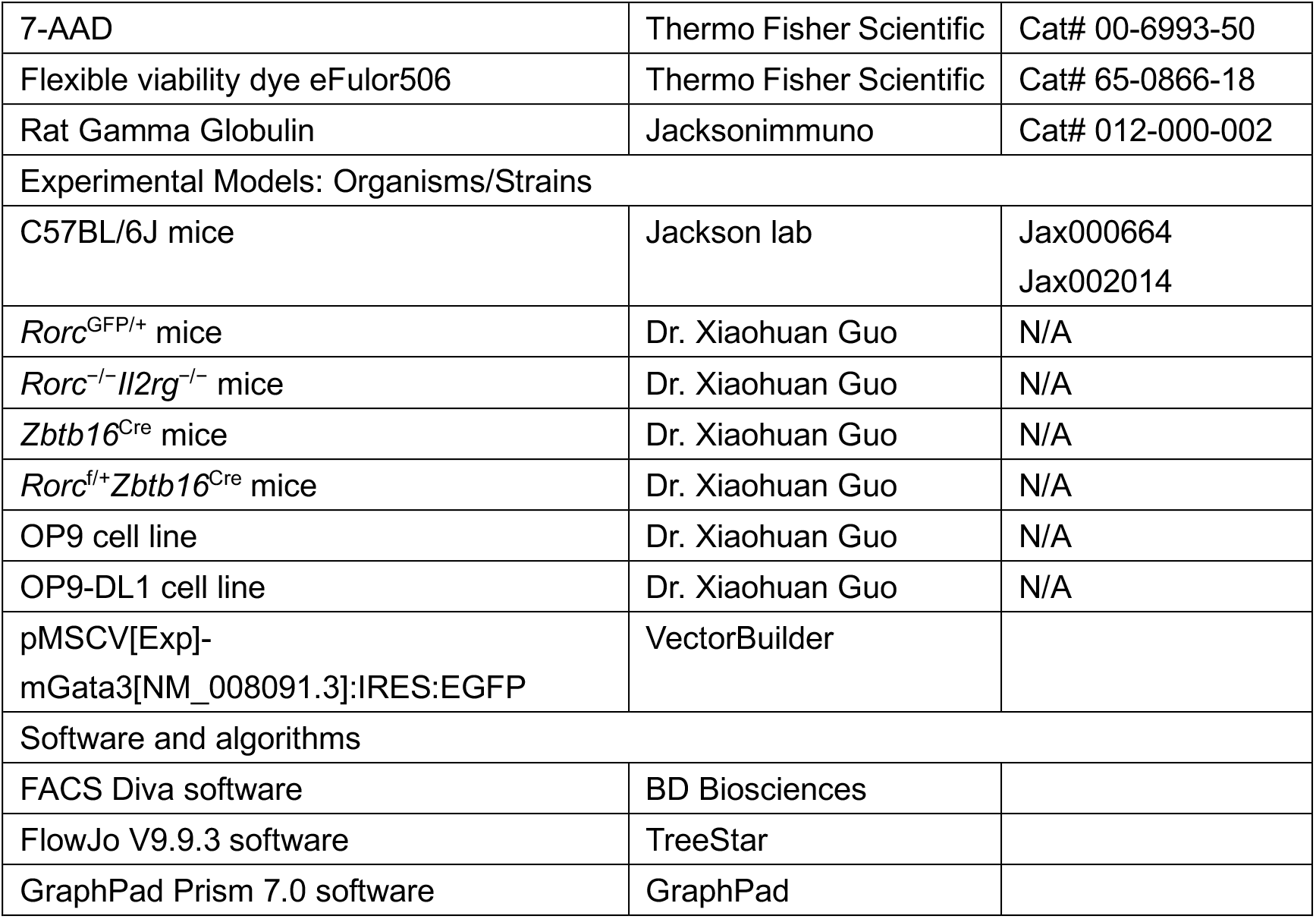

### RESOURCE AVAILABILITY

#### Lead contact

Further information and requests for resources and reagents should be directed to and will be fulfilled by the lead contact, Li Wu.

#### Materials availability

This study did not generate any new unique reagents.

#### Data and code availability

Single-cell RNA-seq data have been deposited at GSA (https://ngdc.cncb.ac.cn/gsa/) with the accession number CRA011319. The codes for scRNA-seq data analysis were available at an online repository (https://github.com/chansigit/ilcdev).

### EXPERIMENTAL MODEL AND SUBJECT DETAILS

#### Animals and ethics statement

CD45.2 mice (Jax000664) and CD45.1 mice (Jax002014) were obtained from Jackson Laboratory. CD45.1.2 mice were generated by crossing CD45.2 mice (Jax000664) and CD45.1 mice (Jax002014). *Rag2*^−/−^*Il2rg*^−/−^ mice, *Rorc*^GFP/+^ mice, *Zbtb16*^Cre^ mice, *Rorc*^f/+^*Zbtb16*^Cre^ mice were kindly provided by Dr. Xiaohuan Guo (Tsinghua University, Beijing, China). Mouse embryos and pups were obtained by in-house breeding. Age of mice were illustrated in indicated experiment. Fetal liver were collected at embryonic day 15.5 (E15.5) or E16.5. For adult bone marrow analysis, 8- to 12-week-old mice were used. Mice after two weeks old were sex-matched, while the male and female fetus or newborn mice were randomly used. All mice used in this study were bred on a C57BL/6 background, and maintained in a specific pathogen free (SPF) animal facility (12 h light and 12 h dark) at Tsinghua University. Animal procedures were under the approval of the Institution Animal Care and Use Committee at Tsinghua University.

### METHOD DETAILS

#### Cell preparation

Single-cell suspensions of bone marrow, fetal liver, small intestine, colon, lung, spleen and liver were prepared following the listed protocols for flow cytometry analysis and/or cell sorting.

For progenitor enrichment from bone marrow, procedures were detailly described elsewhere (Manz et al., 2001; Wu et al., 2001). Briefly, bone marrow cells were enriched by red blood cell removing, density gradient centrifugation (1.086 g/ml Nycodenz), and immunomagnetic depletion of the lineage positive cells labeled by markers of CD3, CD2, CD8, CD19, B220, Mac-1, Ly6G and Ter119. For cell isolation from fetal liver, fetal liver was collected from E15.5 or E16.5 fetus and repeatedly blew by 1 ml pipette tip, cell suspensions were passed through a 70 μM strainer, further enriched by red cell removing and density gradient centrifugation (1.086 g/ml Nycodenz), cells that lighter than 1.086 g/ml were collected and washed. These enriched cells were blocked by Rat IgG and further stained with indicated antibody panel for CLPs (Lin^−^CD127^+^c-Kit^int/+^Flt3^+^α_4_β_7_^−^), αLP1s (Lin^−^CD127^+^c-Kit^int/+^Flt3^+^α_4_β_7_^+^), αLP2s (Lin^−^CD127^+^c-Kit^int/+^Flt3^−^α_4_β_7_^+^).

For cell isolation from the small intestine and colon, the small intestine and colon were flushed clean with cold mouse tissue PBS, opened along the longitudinal axis, and then removed the Peyer’s patches. The residual intestinal tissues were subjected to two sequential 20-min incubated in 20 ml Buffer I (HBSS with 20mM EDTA, 10mM HEPES and 1M DTT), at 37℃ with shaking (220 rpm), for the preliminary digestion. Peyer’s patches and mesenteric tissues were removed from the residual intestinal tissues that were further incubated in 10 ml Buffer II (1640 RPMI + 10% FBS + 1mg/ml Collagenase III) for 60min at 37℃ with shaking (220 rpm), for final digestion. After which, samples were violently vortexed, repeatedly blew by 1 ml pipette tips, and then passed through 70 μM strainer, 40% Percoll density gradient centrifugation was used for further immune cells enrichment. The cells that heavier than 40% Percoll were washed and counted for further analysis.

For lung cell isolation, mouse lung tissues were minced and digested with collagenase 1A (0.5 mg/ml, Sigma) and Dnase I (0.05 mg/ml, Roche) in HBSS for 60 minutes at 37℃ in the thermostatic incubator, with several interval vortexes. Digested lung tissues were further filtrated through a 70 μM strainer, the erythrocytes were removed by using red cell removal buffer (0.168M NH_4_CL), the residual cells were washed and collected for further analysis.

For liver cell isolation from mice after birth, livers were collected and passed through a 100 μM strainer. The cells then enriched by density gradient centrifugation with 40% Percoll, and red blood cells were removed. Splenocytes were isolated by forcing the tissues through a 100 μM strainer, and red blood cells were further removed.

#### Flow cytometry analysis and cell sorting

Flow cytometry staining were performed in 96-well microtiter plates or 5 ml FACS tube. Cells were blocked by Rat IgG and stained with fluorochrome-conjugated antibodies in dark for 30-40 min on ice. Fixable Viability Dye eFluor506 or 7-AAD were used to exclude the dead cells in all these samples. For intracellular transcription factor staining, cells were fixed and permeabilized and stained with indicated antibodies using the Foxp3/Transcription Factor Staining Buffer Set (Thermo Fisher Scientific). Data were acquired on a BD LSRII or Fortessa cytometer, and cells were sorted by FACSAria III, in the flow cytometry core at IITU.

#### *In vitro* differentiation assay of ILC progenitors

We cultured bone marrow progenitors (0.5-1×10^4^/sample) from CD45.1 or CD45.2 wild type mice, including CLP, αLP1, and αLP2, with total bone marrow cells (1×10^4^/sample) from CD45.1.2 wild type or CD45.2 *Rag2^−/−^Il2rg^−/−^*mice as feeder cells in RPMI 1640 media supplemented with 10% fetal bovine serum, glutamine, β-mercaptoethanol, penicillin, and streptomycin. For the OP9 and OP9-DL1 co-culture system, we sorted ILC progenitors (500-1000 cells) and seeded them in wells with OP9 or OP9-DL1 cell lines (500 cells for each well). All cultures were supplemented with stem cell factor (SCF, 100 ng/ml) and IL-7 (20 ng/ml), and ILC subsets among the daughter cells were analyzed 7-8 days later by flow cytometry.

#### Adoptive transfer of ILC progenitors for in vivo differentiation assay

We sorted equal number (0.5∼1×10^4^ for one recipient) of indicated progenitor from fetal liver (CD45.2 wild type) and adult bone marrow (CD45.1,wild type mice), and mixed them with competitor cells (5×10^5^ for one recipient, total bone marrow from CD45.1.2 wild type or CD45.2 *Rag*2^−/−^*Il2rg*^−/−^ mice) befor transferring them to the lethally irradiated mice (CD45.1.2 wild type or CD45.2 *Rag*2^−/−^*Il2rg*^−/−^ mice, 5.5 Gy twice with a 3-hour interval). Neomycin (1 mg/ml) and Polymycin (0.1 mg/ml) were added to the drinking water for 1∼2 weeks.

For the *Rorc* over-expression experiments, we isolated adult bone marrow progenitors (Lin^−^c-Kit^int/+^Sca-1^+^) from wild type (CD45.1) or *Rorc*^Gfp/+^ (CD45.2) mice were transduced them with Rorc-GFP plasmid (*Rorc*^OE^) or empty plasmid (*Rorc*^NC^). For *Gata3* over-expression experiment, we transduced progenitors from the fetal liver (Lin^−^c-Kit^int/+^Sca-1^+^) with either Gata3-GFP plasmid (*Gata3*^OE^) or empty plasmid (*Gata3*^NC^). After Successfully transduction, we sorted and mixed the progenitors with competitor cells (5×10^5^ for each recipient mice, total bone marrow from CD45.2 *Rag*2^−/−^*Il2rg*^−/−^ mice) before transferring to the lethally irradiated mice (CD45.1.2 wild type, 5.5 Gy twice with a 3-hour interval). Neomycin (1 mg/ml) and Polymycin (0.1 mg/ml) were added to the drinking water for 1∼2 weeks. We analyzed cells in the recipient mice 5∼6 weeks after transfer.

#### Single-cell RNA-seq library preparation

Single αLP cells were sorted by FACS from embryonic liver (E15.5∼16.5) and adult bone marrow. After the confirming of cell purity (above 95%) and the cell viability (above 85%) after sorting, cells were further sent to 10×Genomics sequencing platform for single-cell transcriptome sequencing (Annaroad Gene Technology(Beijing)Co., Ltd).

#### scRNA-seq data preprocessing and quality control

The transcriptional profiles of individual cells were acquired through a process involving read mapping, removal of background noises, removal of multiplets, and selection of cells of interest. We mapped scRNA-seq reads to the mm10 version *Mus musculus* reference genome from 10x Genomics (https://cf.10xgenomics.com/supp/cell-exp/refdata-gex-mm10-2020-A.tar.gz) using CellRanger v3.1.0 (10x Genomics) ‘count’ commands. To remove the ambient RNA induced background noises, we removed the used SoupX v1.2.2 (Young and Behjati, 2020) to produce clean UMI-droplet matrices. The clean droplets with low UMI counts and high percentages of mitochondrial expressions were removed from the dataset. Besides single cells, the dataset also contained a small fraction of multiplet droplets. We used DoubletFinder v2.0 (McGinnis et al., 2019) to identify these multiplets. After removing multples, the datasets still contained cells we are not interested in because the cell sorting procedure might have false detections. After data normalization and clustering, we removed the clusters with low expression levels of *Ptprc* (non-immune cells mingled in), clusters with high *Cd3e* expressions (T cells mingled in), clusters with high *Cd79a* expressions (B cells mingled in). The feature selection, normalization, scaling, dimensionality reduction, and visualization of data were conducted with the help of single-cell data analyzing suites Seurat v3.2 (Stuart et al., 2019) and Scanpy v1.7 (Wolf et al., 2018).

To reveal the identities and intrinsic diffenrentiation potential of the hematopoeitic progenitors, we performed cell mapping analysis to show the affinities between the progenitors and other known cell types. We used SingleR v1.0 (Aran et al., 2019) to map the transcriptional profiles in the ImmGen references to our datasets.

#### scRNA-seq data integratoin

To make cell across different developmental stages and batches comparable, we performed integration analyses using a GPU accelerated implementation of the Harmony algorithm, harmony-pytroch v0.1.7 (https://pypi.org/project/harmony-pytorch) (Korsunsky et al., 2019). The analyses reduced unwanted batch-associated variations and produced harmonized cell latent vectors and under the following parameter settings: theta=5, lambda=0.5.

#### Cell vector field analysis

The cell state transition streamlines were derived by RNA velocity analyses (La Manno et al., 2018). Velocyto v0.17 was used to convert the bam format read alignment file produced by CellRanger to the unspliced/spliced counts (La Manno et al., 2018). The cell vector field streamlines and the differential geometry analyses were obtained using Dynamo v1.1.0 (Qiu et al., 2022).

To obtain the representative cells along the velocity trajectories, we calculated the least action path (LAP) along the vector field using the Dynamo package’s dyn.pd.least_action() function. The starting and ending states of the LAPs are selected with the help of the Dynamo vector field’s fixed points. The outputs of LAP analysis contained *T* UMAP points arranged in the state transition’s chronological order. We chose the 10 nearest neighbors in UMAP space for each point and take their average gene expression as a smoothed expression profiles of each point. The cells on the LAP and their neighbors are finally selected as the representative cells along the velocity trajectories. The average gene expressions of each time point creates a representative expression time series (a matrix with *g* rows and *T* columns, *g* is the number of genes in the dataset), which was used for further gene regulatory program analyses.

#### Gene regulatory module identification

Existing methods for identifying gene regulatory program often suffers from compactness problems, where the identified gene expression events are not concentrated around a specific time point. Here, we proposed the TSC (temporal spatial consensus) algorithm to refine the SCENIC gene regulatory network (GRN) output. TSC algorithm combines the spatial localization information (TF-target genes relationships) and the temporal information (gene expression time series). We used the “grn” and “ctx” command in pySCENIC v0.11 (Van de Sande et al., 2020) to generate a basic GRN backbone.

TSC accepted the LAP derived gene expression time series (*g* x *T* matrix) and calculated dynamic time wrapping distances (DTW) between every gene’s expression time series (https://zenodo.org/records/7158824). A k-NN (k nearest neighbor) graph was then created using these distances (k=20 by default) and edges on the graph were next dropped out if the two connecting nodes were not mutual k-NN. Given the DTW graph and the SCENIC graph as two layers sharing the same set of nodes (genes), we build a multiplex network. We used Node2Vec (Grover and Leskovec, 2016) for layers of the multiplex network and obtained latent embedding for each genes. A hierarchical clustering from the scipy.cluster.hierarchy package was used to group these genes into different affinity modules, which was the TSC-refined gene regulatory modules.

#### Differentially Expressed Genes and GSVA analysis

The DEG (differentially expressed genes) were identified with the Seurat package’s FindMarkers() or FindAllMarkers() function with an adjusted p-value <0.05 threshold. The DEG analyses compared expressions using Wilcoxon rank sum test on the normalized data slot in Seurat objects.

In Gene Set Variation Analysis (GSVA) (Hanzelmann et al., 2013), cells are firstly overclustered within cell types to be analyzed, and the overclustered cells are then merged into metacells by taking their average expression profiles. GSVA analyses were performed on these metacells using gene sets from MSigDB Hallmark (Liberzon et al., 2011).

We used the ternary plot (Hamilton and Ferry, 2018) to present the tangled relationships between the *Gata3* regulon, the *Rorc* regulon, and the Notch signaling. The *Gata3* and *Rorc* regulons’ gene members were extracted from the SCENIC-derived gene regulatory modules. We used the following Notch signaling pathway gene members: *Notch1*, *Notch2*, *Notch3*, *Notch4*, *Furin*, *Psen2*, *Adam17*, *Dtx1*, *Dtx4*, *Rbpj*. The activation scores of the three modules were evaluated by GSVA. The samples used for GSVA analysis were the average expression profiles of the time steps derived from the LAP analyses.

### QUANTIFICATION AND STATISTICAL ANALYSIS

#### Statistical Analysis

GraphPad Prism 7 was used for statistical analysis with the specific tests and the number of samples used for each analysis (n) shown in the figure legends.

---

**Figure S1.**
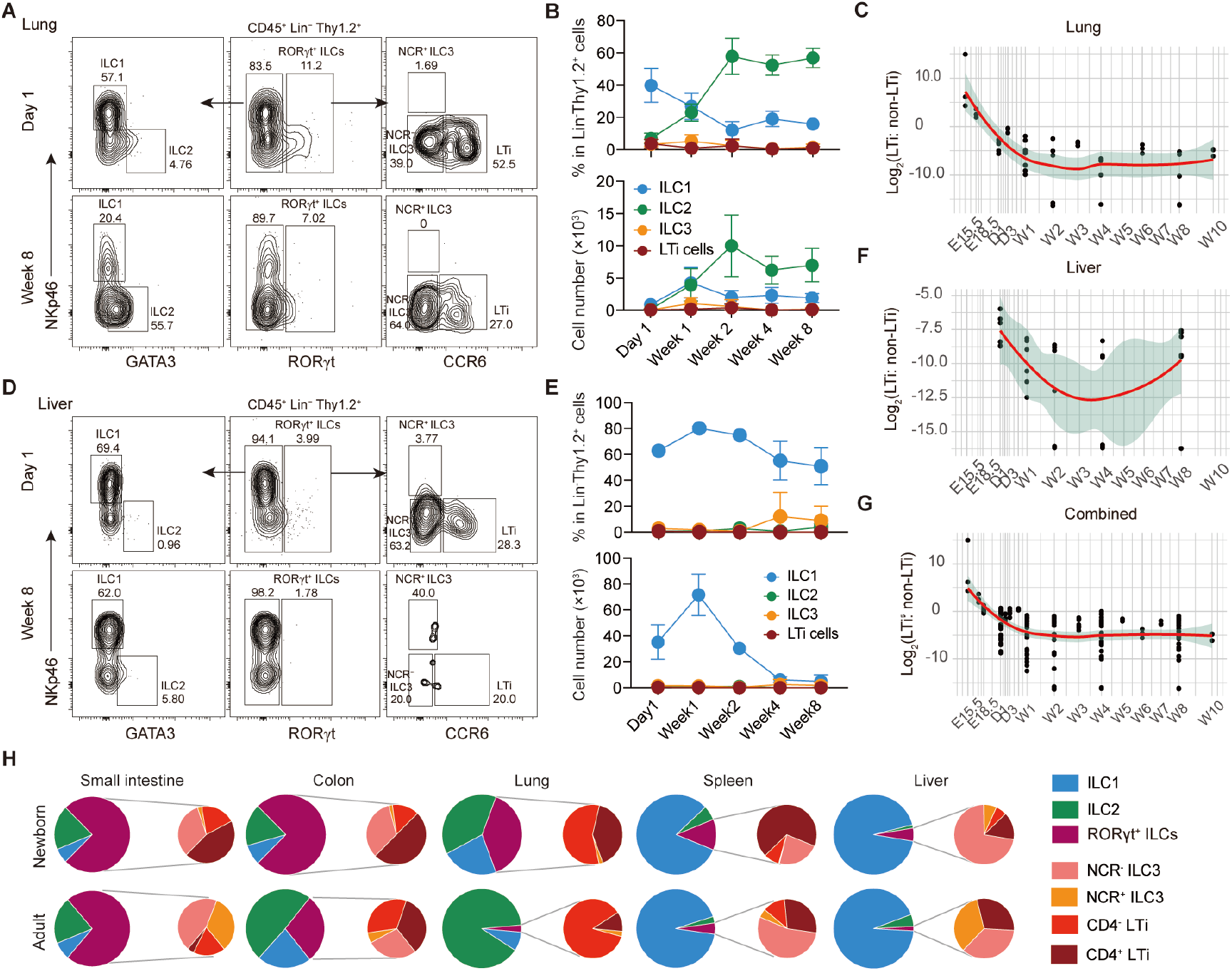
ILC Subsets in lung and liver during ontogeny. Related to Figure 1. (A and D) Representative FACS plots display the frequencies of major ILC subsets in (A) lung and (D) liver of the newborn (Day 1) and adult (Week 8) mice. (B and E) Percentage and absolute number of ILC subsets in (B) lung and (E) liver during ontogeny are presented. Data (n=3 mice for each group) represent two independent experiments and are shown as mean ± SD. (C, F and G) The logarithm of the percentage ratio of LTi cells to non-LTi ILCs in (H) liver, (I) lung, and (G) all these tissues combined. Each dot represents the value for one individual mouse. (H) A pie graph is provided to illustrate the composition of ILC subsets across tissues between the newborn (Day 1) and adult (Week 8).

**Figure S2.**
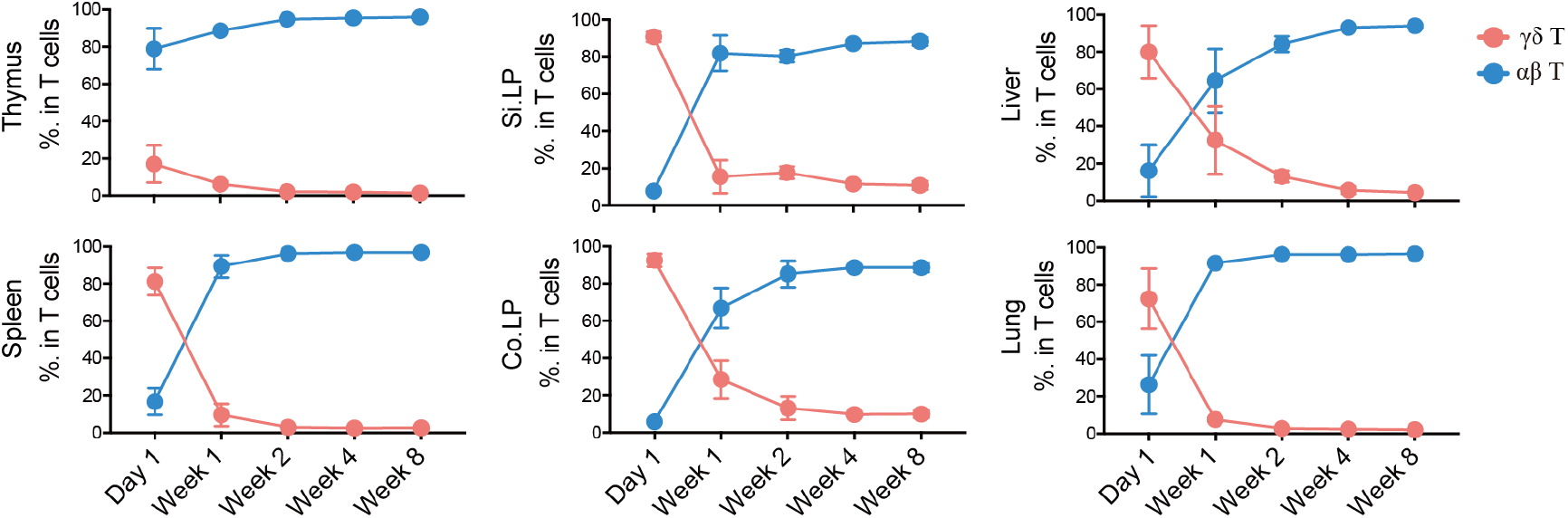
Ontogeny of αβ T cells and γδ T cells. Related to Figure 1. The percentage of the αβ T cells and γδ T cells in indicated tissues during ontogeny. Data (n= 6 mice per group) are pooled from two independent experiments (mean ± SD).

**Figure S3.**
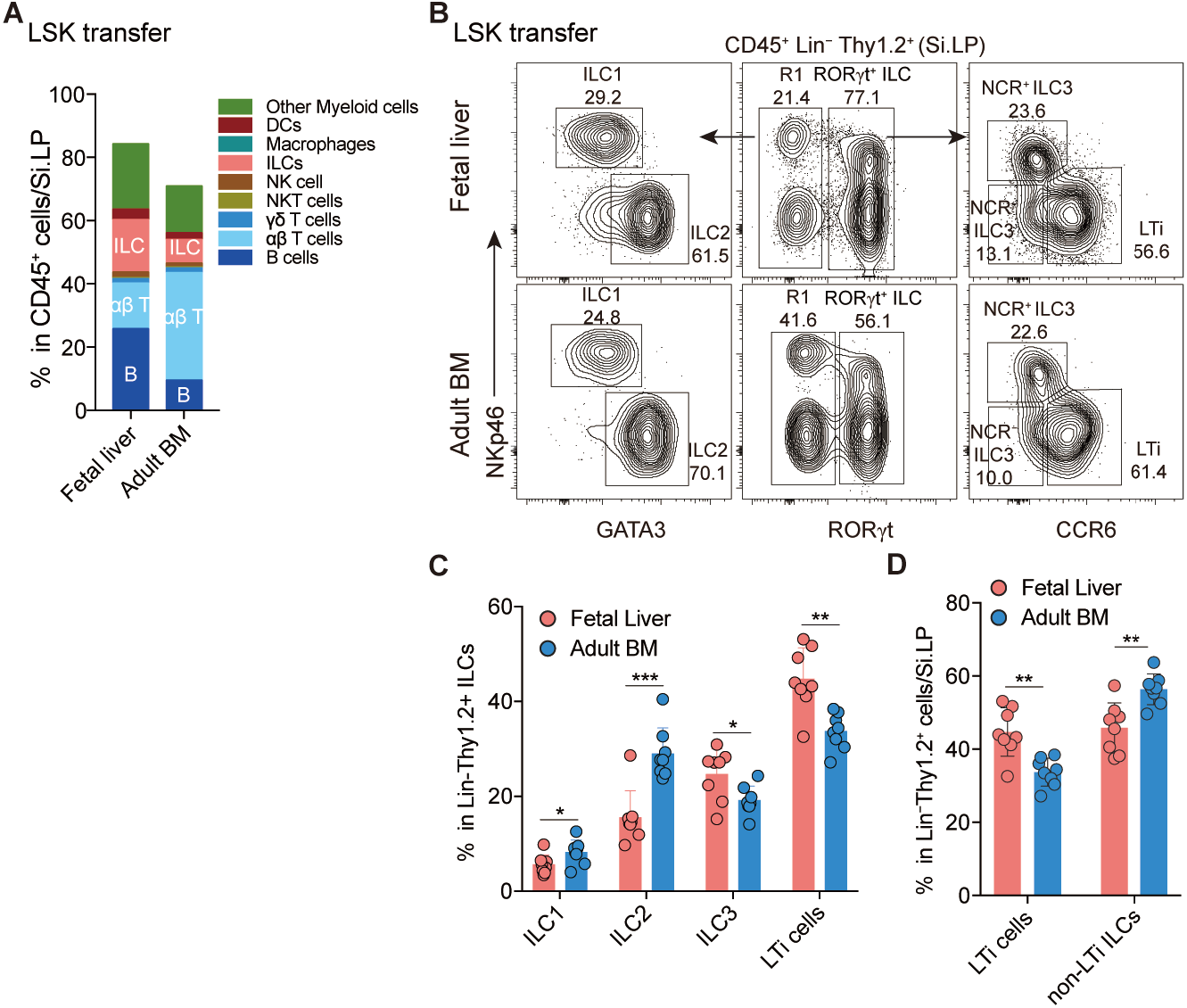
ILC subsets generated by fetal and adult hematopoietic progenitors. Related to Figure 2. (A) Composition of major immune cells derived from LSKs of fetal liver and adult bone marrow in the recipient small intestine. Data (n=8 mice pre group) are pooled from three independent experiments. (B) Representative FACS plots show the major ILC subsets derived from LSKs of fetal liver and adult bone marrow in the recipient small intestine. (C) Percentage of major ILC subsets derived from fetal and adult LSKs in the recipient small intestine. (D) Proportion of LTi cells and non-LTi ILCs derived from LSKs of fetal liver and adult bone marrow in the recipient small intestine. Data (n= 8 mice per group) are pooled from three independent experiments.

**Figure S4.**
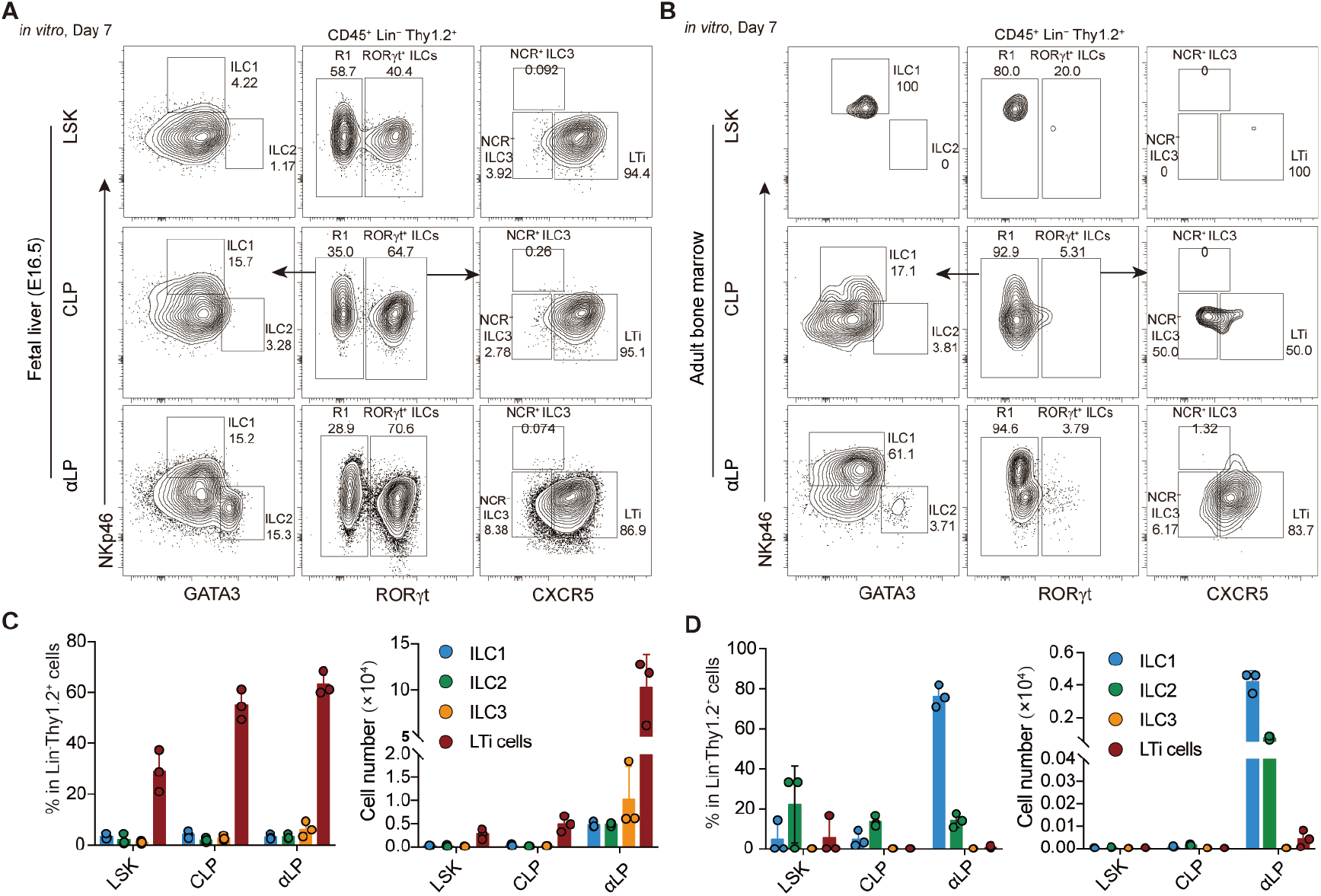
Differentiation assay of ILC progenitors from fetal liver and adult bone marrow. Related to Figure 2. (A-B) Representative FACS plots display the ILC subsets derived from (A) fetal liver and (B) adult bone marrow LSKs, CLPs and αLPs, after seven days of culture supplemented with IL-7 and SCF *in vitro*. (C-D) The composition of major ILC subsets among the daughter cells derived from the indicated counterpart progenitors from (C) fetal liver and (D) adult bone marrow are presented. Data (n=3 samples per group) represent two independent experiments and are shown as mean ± SD.

**Figure S5.**
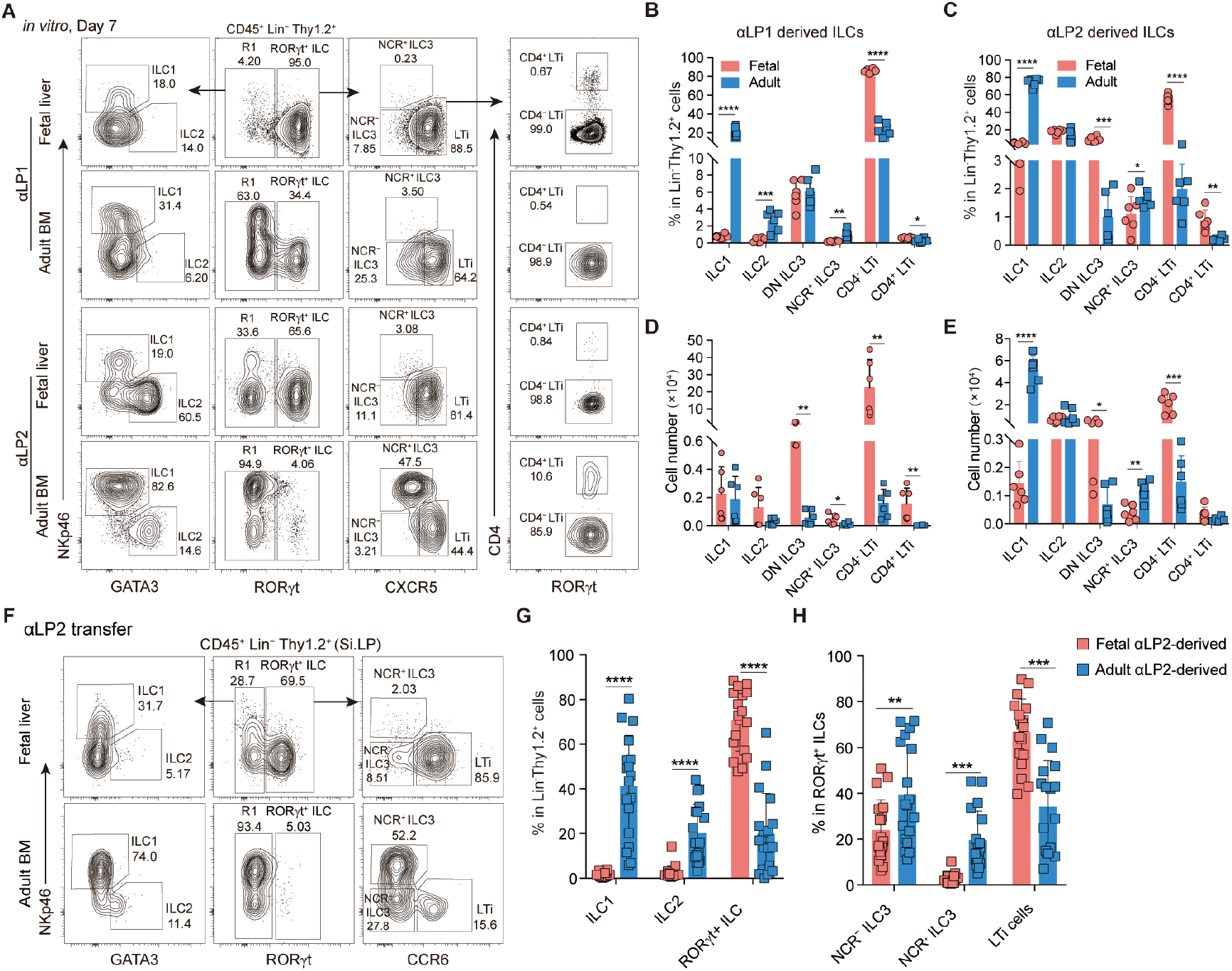
Differentiation assay of αLP1s and αLP2s from fetal liver and adult bone marrow. Related to Figure 2. (A) Representative FACS plots display the ILC subsets derived from fetal and adult αLP1s or αLP2s after seven days of culture with IL-7 and SCF *in vitro*. (B-E) The (B-C) percentage and (D-E) absolute number of major ILC subsets derived from fetal and adult (B, D) αLP1s or (C, E) αLP2s are presented. Data (n=6 samples per group) are pooled from two independent experiments and are shown as mean ± SD. (F) Representative FACS plots show ILC subsets derived from fetal and adult αLP2s in the small intestine of recipient mice. (G-H) Percentage of (G) ILC1, ILC2 and RORγt^+^ ILCs and (H) RORγt^+^ ILC subsets derived from fetal and adult αLP2s in the recipient small intestine. Data (n=18 samples per group) are pooled from seven independent experiments (mean ± SD).

**Figure S6.**
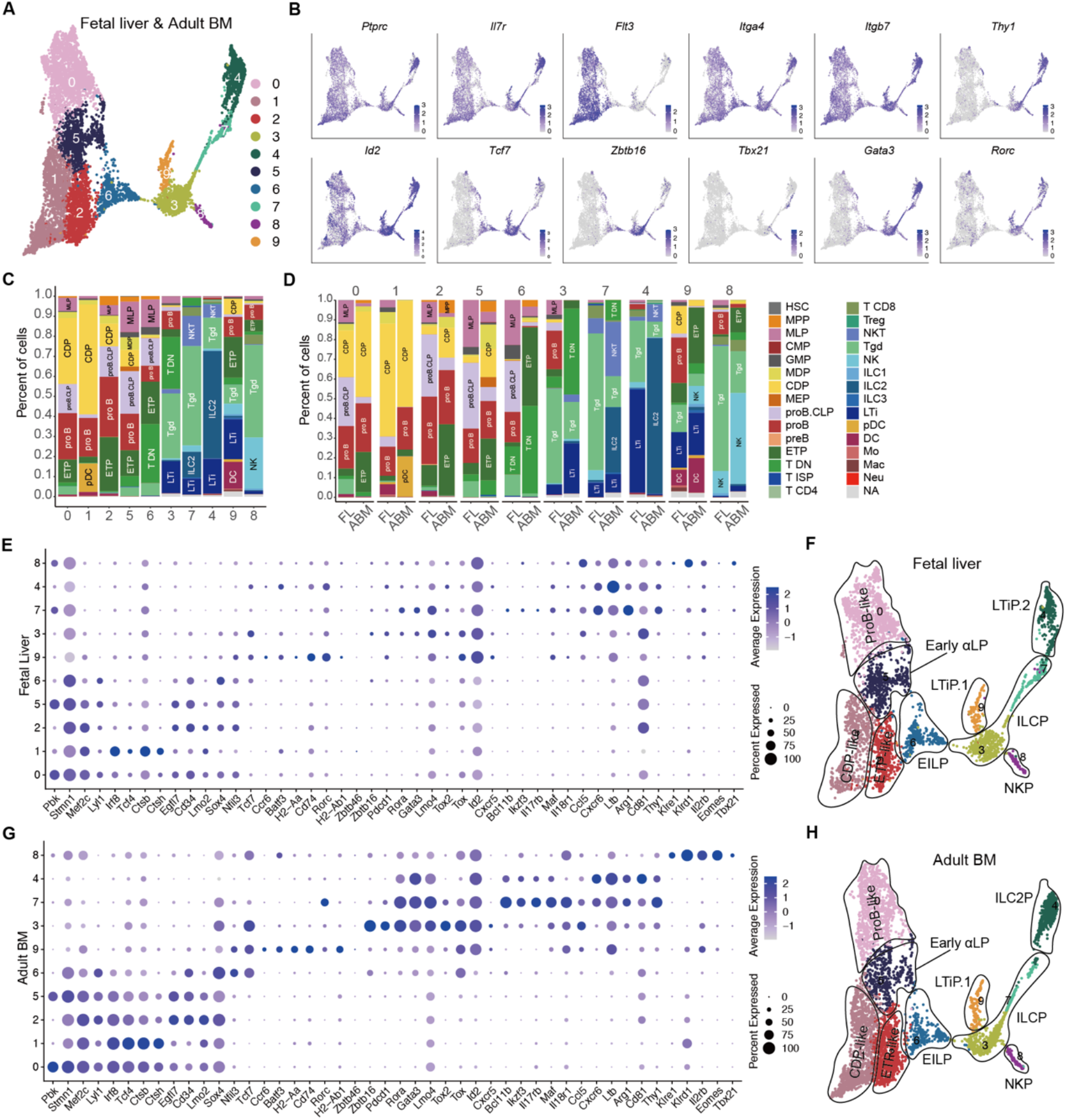
Single-cell transcriptomic landscape of fetal and adult ILC progenitors. Related to Figure 3. (A) The cellular composition of αLPs from integrated fetal liver and adult bone marrow is visualized using Uniform Manifold Approximation and Projection (UMAP). Cell types are color-coded. (B) Marker gene expression is shown across the identified clusters among αLPs by feature-plot analysis. (C and D) Histograms show the top correlated cell types in each (C) integrated or (D) separated cluster by comparing the single-cell transcriptome to that in the ImmGen database. (E and G) Dot-plot representations of the marker gene expression for the identified clusters among αLPs from (E) fetal liver and (G) adult bone marrow are presented. (F and H) The indicated clusters in (F) fetal liver and (H) adult bone marrow αLPs are finely annotated.

**Figure S7.**
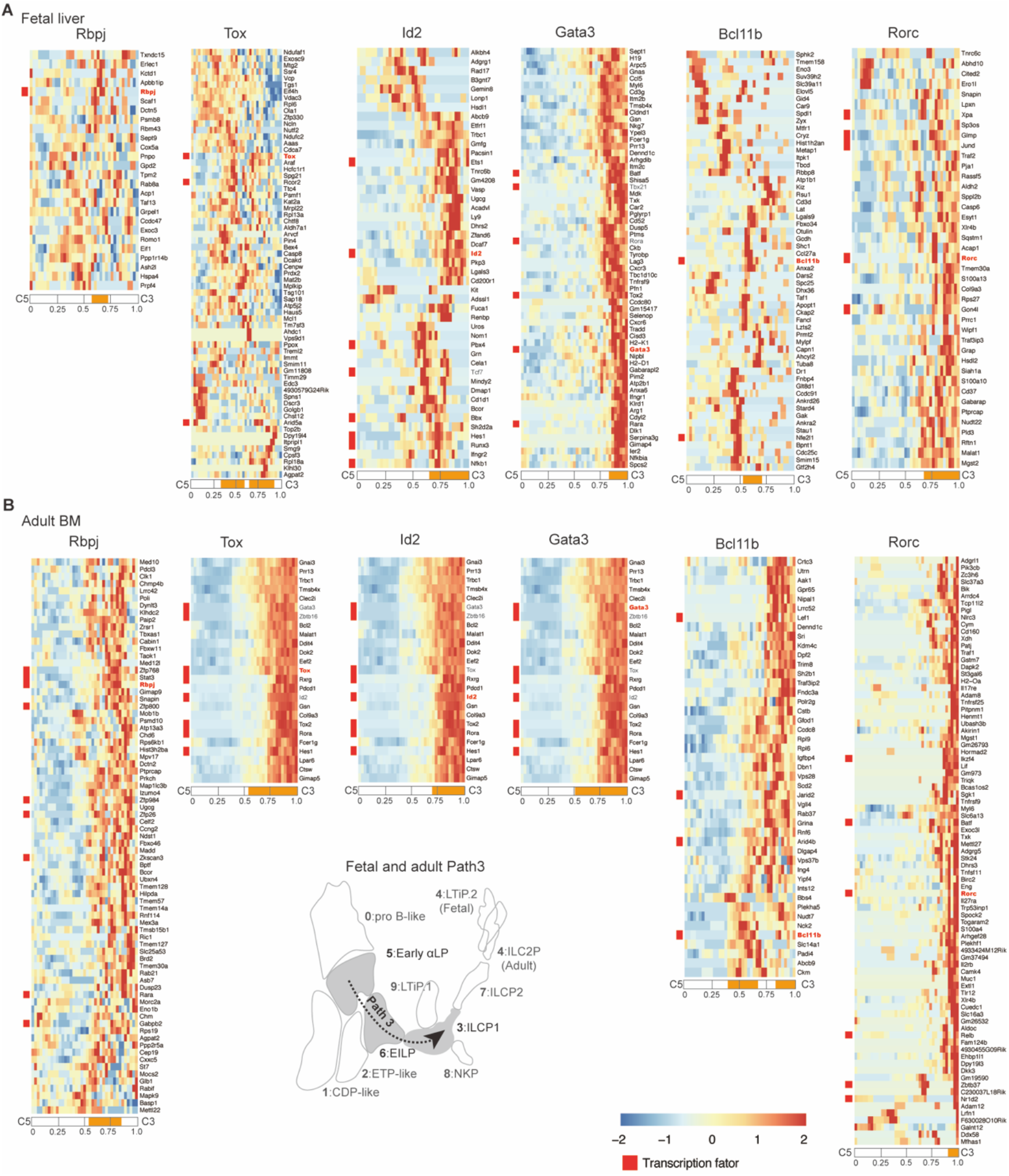
TSC analysis of indicated transcription factors during fetal and adult ILC development revealed by single-cell RNA-sequence. Related to Figure 4. (A and B) Heatmap showing the expression of genes accompanied with indicated transcription factors along the ILC differentiation pseudotime (path 3) predicted by TSC analysis in (A) fetal liver and (B) adult bone marrow. The red characters mark the transcription factors. The pseudotime from C5 to C3 (path 3) is divided into four equal parts. The orange characters bend during the pseudotime, representing the expression period of the indicated transcription factor.

**Figure S8.**
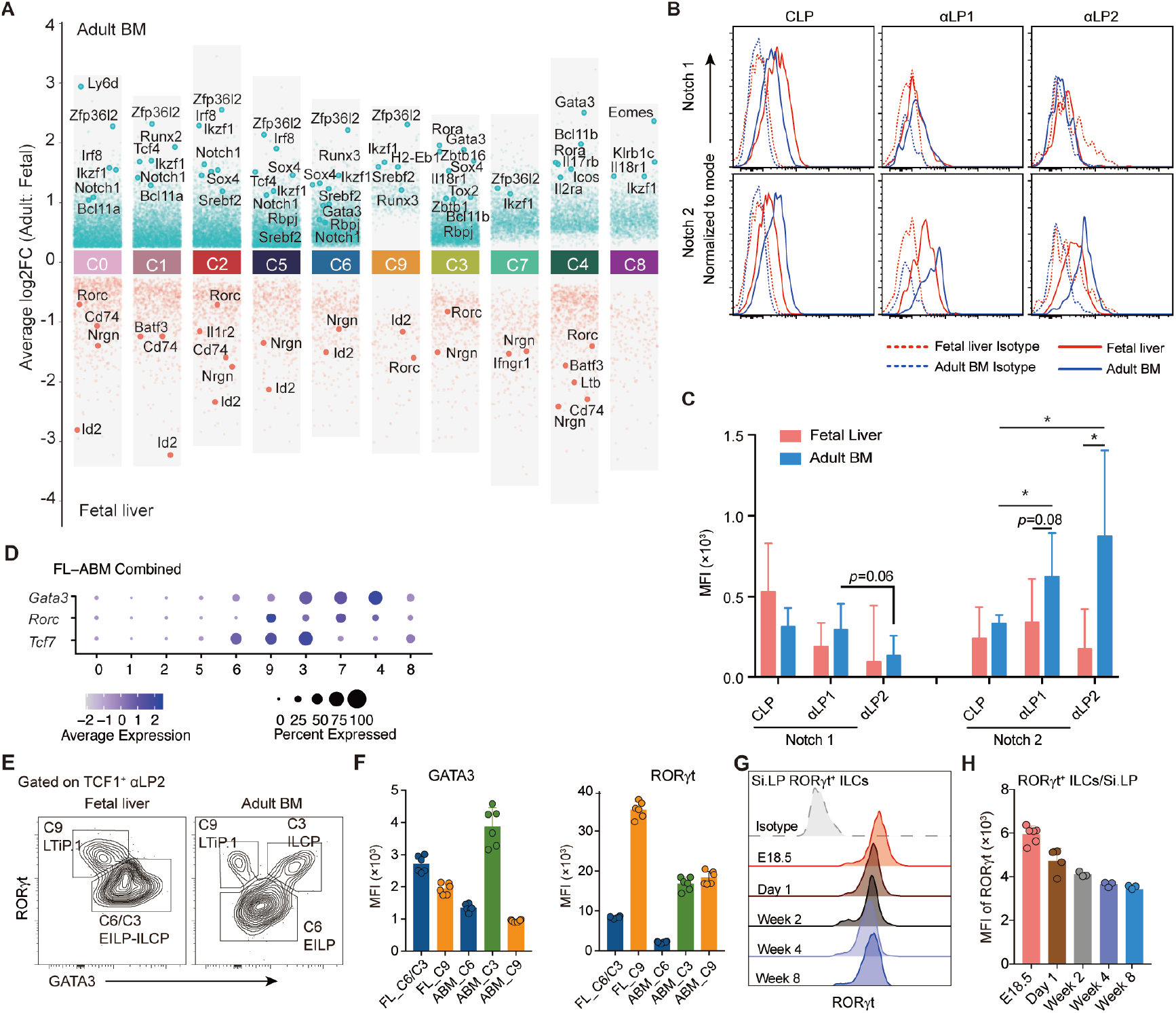
Different expression of genes involved in Notch signaling, *Gata3* and *Rorc* between fetal and adult. Related to Figure 4. (A) Differentially expressed genes between indicated clusters in fetal liver and adult bone marrow. (B and C) Flow cytometry analysis showing the expression of Notch1 and Notch2 on ILC progenitors, including CLPs, αLP1s and αLP2s, from both fetal liver and adult bone marrow. Data (n=6-7 samples per group) are pooled from three independent experiments and are presented as mean ± SD. (D) Dot plots showing expression features of *Tcf7*, *Gata3*, and *Rorc* in ILC progenitors. (E-F) Flow cytometry analysis showing expression of RORγt and GATA3 in fetal and adult TCF1^+^ αLP2s. Data (n=6 per group) represent two independent experiments (mean ± SD). (G) Flow cytometry plots showing the intracellular expression level of RORγt in intestinal RORγt^+^ ILCs at different stages. (H) Mean fluorescence intensity (MFI) of RORγt in small intestinal RORγt^+^ ILCs during ontogeny. Data (n=3-6 per group) are pooled from two independent experiments and shown as mean ± SD.

**Figure S9.**
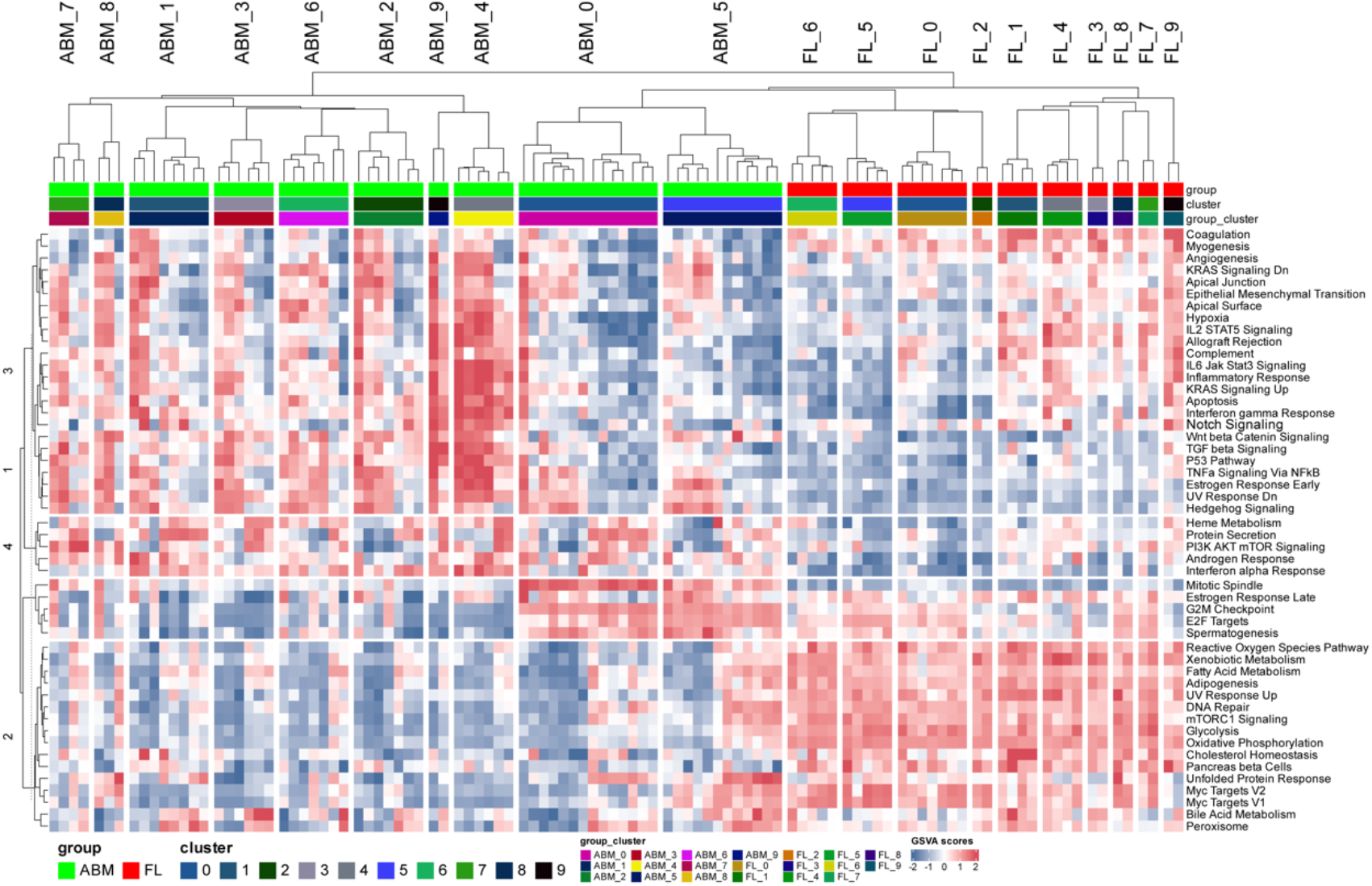
Differentially distributed signaling pathways between fetal and adult αLPs. Related to Figure 4. Gene signature analyses and clustering of gene set variation analysis (GSVA) scores for single αLPs from fetal liver (Right) and adult bone marrow (Left).

**Figure S10.**
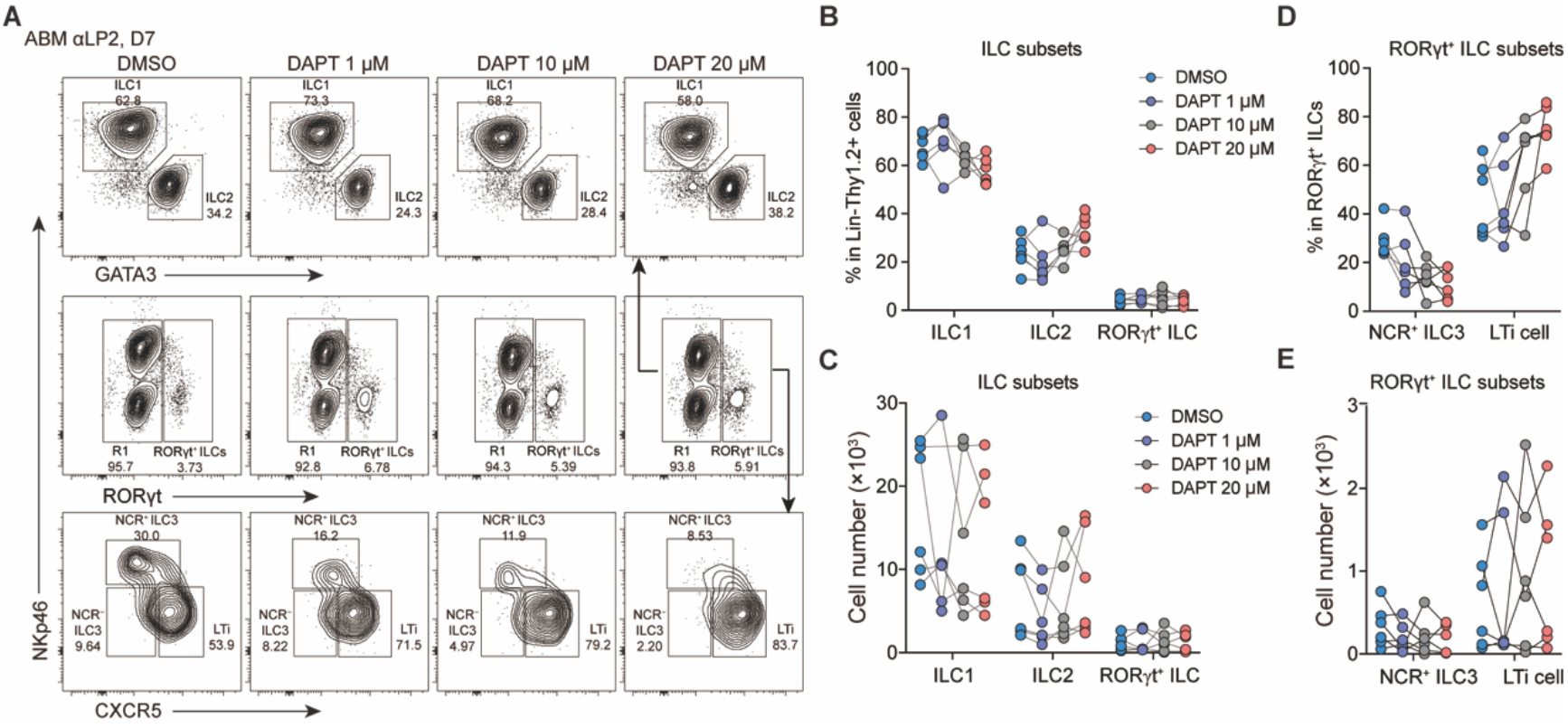
Blocking Notch signaling led to increased LTi cells but decreased NCR^+^ ILC3s generation by adult αLP2s. Related to Figure 5. (A) Notch signaling inhibitor DAPT was administered during the *in vitro* differentiation of adult bone marrow αLP2s with varying concentrations. Representative flow cytometry plots showing the major ILC subsets among the daughter cells. (B-C) Percentage (B) and cell number (C) of ILC1s, ILC2s, and RORγt^+^ ILCs in (A). (D-E) Percentage (D) and cell number (E) in (A). Data (n=6 per group) represent three independent experiments (mean ± SD).

**Figure S11.**
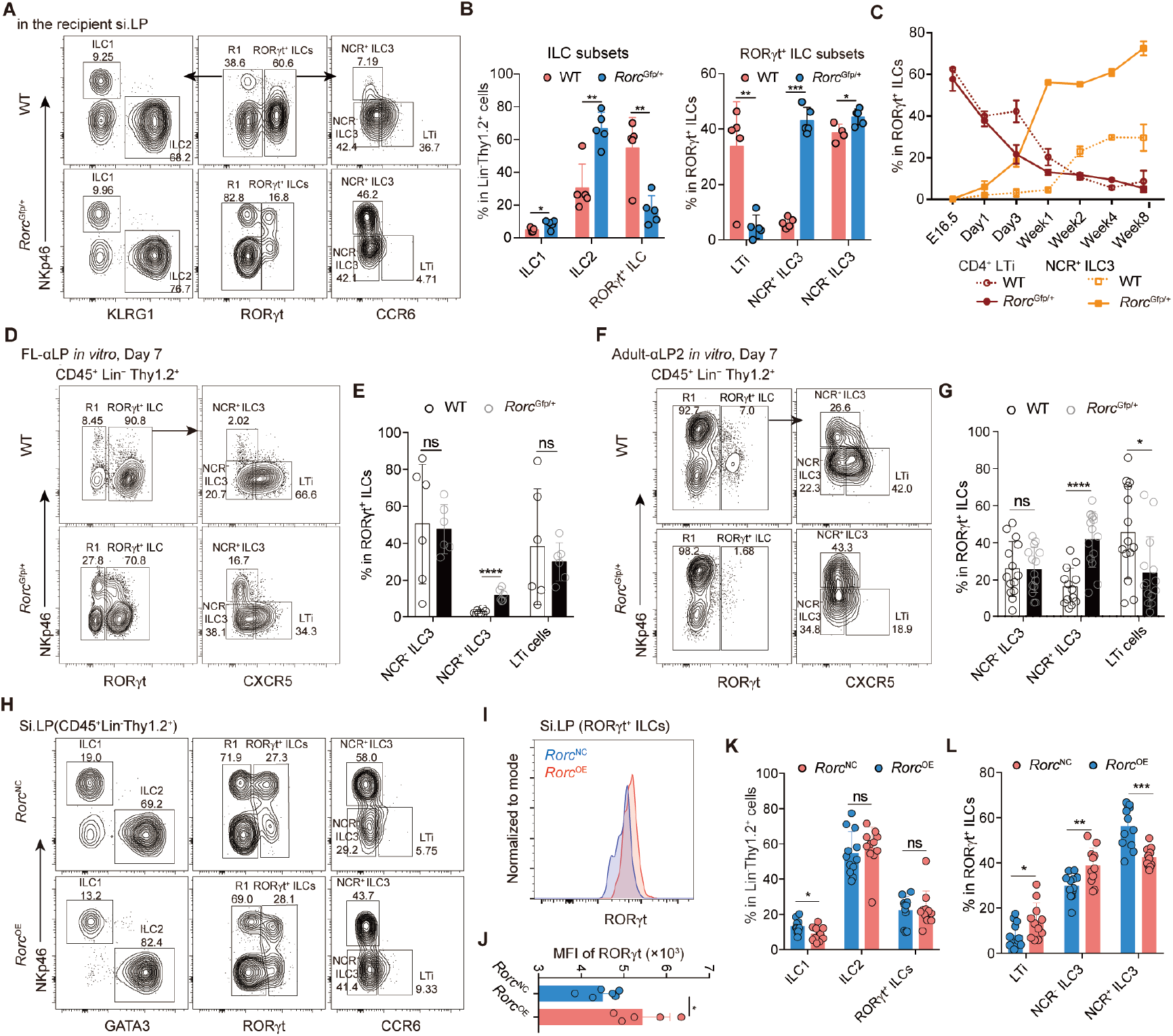
RORγt reduction promotes NCR^+^ ILC3 differentiation. Related to Figure 5. (A) Representative FACS plots show the small intestinal ILC subsets derived from wide type (WT) and *Rorc*^Gfp/+^ bone marrow progenitor (Lin^−^c-Kit^+^ cells) and in the recipient mice. (B) The percentage of major ILC subsets in (A). Data (n=5-6 per group) are pooled from two independent experiments and shown as mean ± SD. (C) Percentage of NCR^+^ ILC3 and CD4^+^ LTi cells in the small intestine of both wide type and *Rorc*^Gfp/+^ mice during ontogeny. (D-G) Fetal liver αLPs or adult bone marrow αLP2s are sorted from wide type and *Rorc*^Gfp/+^ mice for *in vitro* culture supplemented with IL-7 and SCF for 7 days. FACS plots and percentage of RORγt^+^ ILC subsets among the (D and E) fetal liver-derived and (F-G) adult bone marrow-derived daughter cells were shown. Each circle in the graph represented a biological replicate (n=6-14 per group pooled from two to five independent experiments). (H-L) LSK from adult *Rorc*^Gfp/+^ mice were transduced by empty (*Rorc*^NC^) or *Rorc* (*Rorc*^OE^) plasmid. (H) Flow cytometry showing the small intestinal ILC subsets in the recipient mice. (I) Intracellular RORγt staining and (J) indicated MFI in intestinal RORγt^+^ ILCs in (H). (K-L) Percentage of major ILC subsets derived from empty (*Rorc*^NC^) or *Rorc* (*Rorc*^OE^) plasmid-transduced progenitors. Data (n=8-9 per group) are pooled from three independent experiments and shown as mean ± SD.

